# SAM homeostasis is regulated by CFI_m_-mediated splicing of MAT2A

**DOI:** 10.1101/2020.11.12.380626

**Authors:** Anna M. Scarborough, Juliana N. Flaherty, Olga V. Hunter, Kuanqing Liu, Ashwani Kumar, Chao Xing, Benjamin P. Tu, Nicholas K. Conrad

## Abstract

S-adenosylmethionine (SAM) is the methyl donor for nearly all cellular methylation events. Cells regulate intracellular SAM levels through intron detention of the MAT2A RNA, which encodes only SAM synthetase expressed in most cells. The N^6^-adenosine methyltransferase METTL16 promotes splicing of the MAT2A detained intron by an unknown mechanism. Using an unbiased CRISPR knock-out screen, we identified CFI_m_25 (NUDT21) to be a regulator of MAT2A intron detention and intracellular SAM levels. CFI_m_25 is a component of the cleavage factor I_m_ (CFI_m_) complex that regulates poly(A) site selection, but we show it promotes MAT2A splicing, independent of poly(A) site selection. CFI_m_25-mediated MAT2A splicing induction requires the RS domains of its binding partners, CFI_m_68 and CFI_m_59 as well as binding sites in detained intron and 3′ UTR. These studies uncover mechanisms that regulate MAT2A intron detention and reveal previously undescribed roles for CFI_m_ in splicing and SAM metabolism.

## INTRODUCTION

S-adenosylmethionine (SAM) is the methyl donor used as a co-factor for methyltransferases that modify DNA, RNA, and proteins. Due to the widespread use of methylation in the regulation of nearly all cellular functions, SAM may be the second most widely used metabolite next to ATP (Walsh et al., 2018). The *MAT2A* gene encodes MATα2, which is only SAM synthetase expressed in nearly every tissue except liver (Murray et al., 2019). The enzyme produces SAM using ATP and L-methionine, so SAM levels are closely linked with methionine metabolism. The continual turnover of SAM in the cell by methyltransferases and its central importance to a wide variety of cellular pathways requires that its levels be constantly monitored. Indeed, disruption of the methionine-SAM homeostasis has widespread consequences for health and disease (Gao et al., 2019; Lio and Huang, 2020; Ouyang et al., 2020; Parkhitko et al., 2019).

We identified the N^6^-methyladenosyl (m^6^A) transferase METTL16 as a key regulator of SAM-responsive MAT2A RNA splicing and proposed that it serves as a critical intracellular SAM sensor (Pendleton et al., 2017). METTL16 methylates MAT2A on six evolutionarily conserved hairpins (hp1-hp6) in the 3′ untranslated region (3′UTR; Fig 1A) (Parker et al., 2011; Pendleton et al., 2017; Warda et al., 2017). These hairpins contain UACAGARAA (R=G or A) motifs which, in combination with their structure, allows for methylation by METTL16 at the A4 position (Doxtader et al., 2018; Mendel et al., 2018; Pendleton et al., 2017). METTL16 interactions with MAT2A hairpins appear to regulate MAT2A expression by two mechanisms. Upon methylation of hp2-6, the cytoplasmic stability of the MAT2A mRNA decreases (Martinez-Chantar et al., 2003; Pendleton et al., 2017; Shima et al., 2017). This reflects a similar “reader-writer” paradigm typically used for the m6A marks deposited by the primary mRNA methyltransferase complex METTL3-METTL14-WTAP (Meyer and Jaffrey, 2017; Yue et al., 2015).

**Figure 1.**
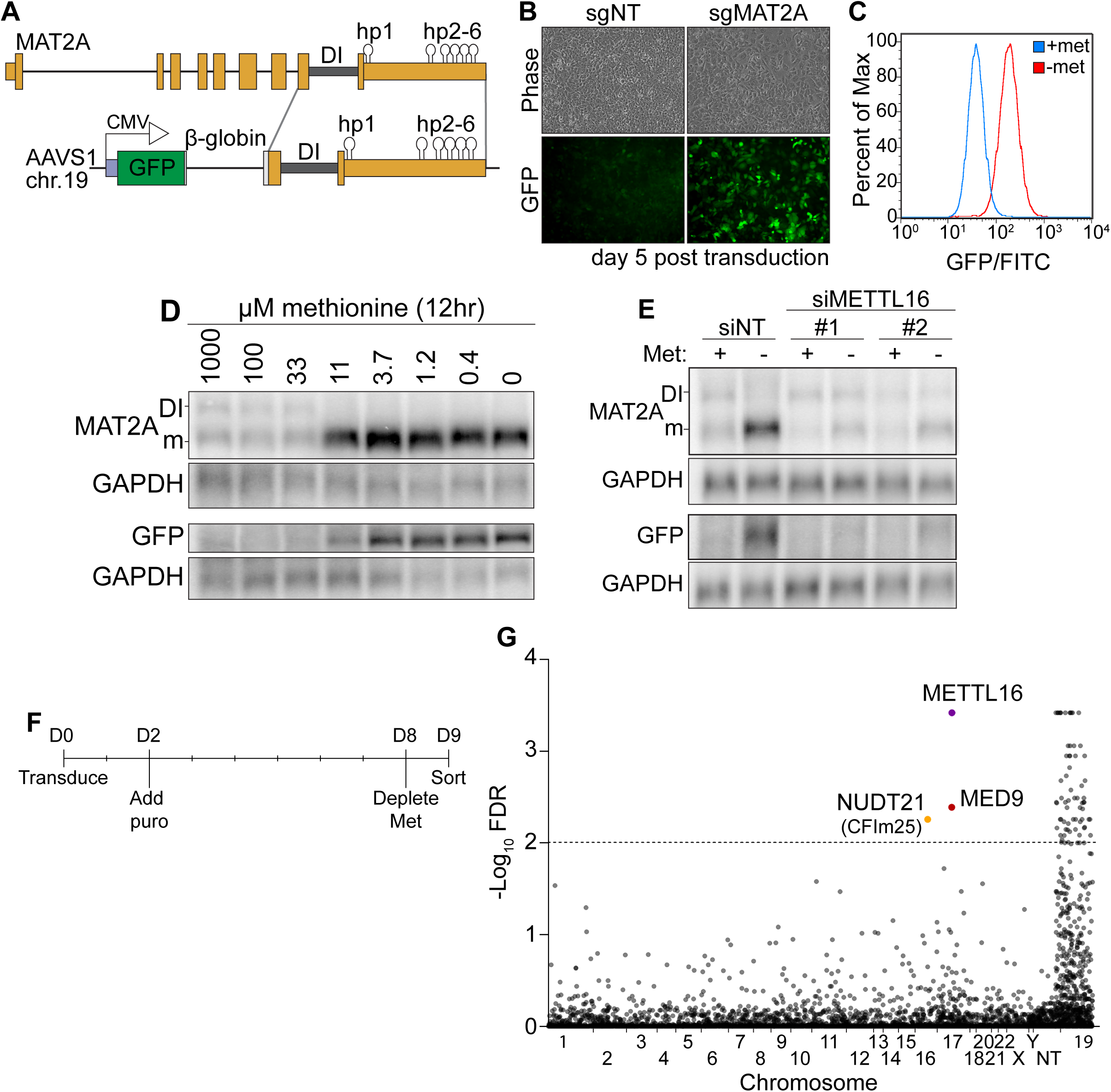
**A CRISPR Screen Identifies CFI_m_25 as a Candidate MAT2A Splicing Factor** A. Diagram of endogenous MAT2A gene (top) and GFP-MAT2A reporter (bottom). DI, detained intron; hp, hairpins; diagram is not to scale. B. Representative images of reporter cells (phase) and GFP five days after transduction with lentivirus expressing Cas9 and sgRNA targeting MAT2A or non-targeting (sgNT) control. Cells were maintained in methionine-rich media. C. Flow cytometry results monitoring GFP production in the reporter cell line after 24-hour conditioning in methionine-rich (blue) or methionine-free (red) media. Displayed as percent of maximum cell count for a given GFP intensity. D. Northern blot analysis of GFP and endogenous MAT2A RNAs produced from the reporter line after 12 hours in media with the indicated methionine concentrations. In all figures, DI and m mark the MAT2A detained intron and mRNA isoforms, respectively. E. Northern blot analysis of GFP and endogenous MAT2A RNA expression after four hours ± methionine depletion in the reporter line. Cells were treated with non-targeting control (siNT) or METTL16 (siMETTL16) siRNAs. F. CRISPR screen timeline. G. CRISPR screen results. CRISPR screen was performed in three biological replicates before analysis by MAGeCK. The -log_10_(FDR) of the analysis is plotted on the y-axis and genes are organized alphabetically by chromosome number on the x-axis. Non-targeting (NT) guides are also included. Genes above the dotted line have an FDR < 0.01.

Here we focus on the second mechanism for METTL16-mediated regulation of MAT2A in which it controls splicing of the last intron of the MAT2A transcript. Our working model proposes that in SAM-rich conditions, METTL16 methylates hp1 then dissociates, resulting in increased retention of the last intron (Fig 1A)(Pendleton et al., 2017; Pendleton et al., 2018). Failure to excise the last intron of MAT2A results in nuclear retention and degradation of the transcript, so this intron is defined as a detained intron (DI)(Boutz et al., 2015; Bresson et al., 2015; Pendleton et al., 2018). Upon SAM limitation, METTL16 binds hp1 but low SAM levels lead to decreased enzymatic turnover of METTL16. The resulting increased METTL16 occupancy on hp1 promotes splicing of the final MAT2A intron. Two carboxyl-terminal vertebrate conserved regions (VCR) in METTL16 are necessary and sufficient to promote splicing of MAT2A reporters and increase the affinity of METTL16 for RNA (Aoyama et al., 2020; Pendleton et al., 2017). Thus, with respect to hp1 and regulation of intron detention, METTL16 functions as both the m6A writer and reader. However, METTL16 lacks recognizable splicing domains and has no known protein interacting partners, so its mechanism for promoting splicing of the MAT2A DI has been unclear (Ignatova et al., 2019).

Splicing of terminal introns is coupled with 3′-end formation. Definition of both the 5′ and 3′ ends of an exon is necessary for splicing, but the lack of downstream introns requires that terminal exons use a different mechanism (De Conti et al., 2013; Martinson, 2011; Niwa et al., 1992; Niwa et al., 1990). In this case, splicing factors and 3′ end formation machinery directly interact during the definition of the terminal exon (Davidson and West, 2013; Dettwiler et al., 2004; Kyburz et al., 2006; Lutz et al., 1996; McCracken et al., 2002; Millevoi et al., 2006; Rappsilber et al., 2002; Shi et al., 2009; Vagner et al., 2000). While splicing of terminal exons and 3′ end formation are reciprocally stimulatory, the mechanisms of this coupling remain incompletely understood.

The cleavage factor I_m_ complex (CFI_m_) is a component of the 3′-end formation machinery with characteristics that suggest it links 3′ end formation with splicing (Hardy and Norbury, 2016; Martinson, 2011). The CFI_m_ complex consists of a dimer of CFI_m_25 (NUDT21, CPSF5) and each CFI_m_25 interacts with a monomer of CFI_m_68 (CPSF6) or CFI_m_59 (CPSF7) to form a heterotetrameric CFI_m_ complex (Kim et al., 2010; Ruegsegger et al., 1996; Ruegsegger et al., 1998). The CFI_m_ complex was identified as a 3′-end formation factor, but more recent work suggests it is not required for polyadenylation of all mRNAs. Instead, CFI_m_ is an enhancer that regulates the efficiency of poly(A) site usage (Zhu et al., 2018). As a result, knockdown of CFI_m_ components leads to widespread changes in polyadenylation, with the predominant effect being a shift to proximal poly(A) site usage (Brumbaugh et al., 2018; Gruber et al., 2012; Li et al., 2015; Martin et al., 2012; Masamha et al., 2014; Zhu et al., 2018). CFI_m_68 and CFI_m_59 contain arginine- and serine-rich regions (RS domains) that aid in 3′ end formation (Zhu et al., 2018). However, RS domains are common components of splicing factors, suggesting a role for CFI_m_68 and CFI_m_59 in splicing (Dettwiler et al., 2004; Hardy and Norbury, 2016; Long and Caceres, 2009; Millevoi et al., 2006; Rappsilber et al., 2002; Ruegsegger et al., 1998). Thus, in addition to its well-defined roles in 3′-end formation and alternative polyadenylation (APA), the CFI_m_ complex may function in splicing. To date, there has been little direct evidence that any specific pre-mRNA requires CFI_m_ as a splicing factor independent of its functions in poly(A) site selection.

Here, we uncover the role of the CFI_m_ complex in METTL16-mediated splicing of MAT2A. Using a SAM-sensitive GFP reporter in a CRISPR knock-out screen, we identified CFI_m_25 as a candidate factor in the regulation of MAT2A intron detention. Similar to loss of METTL16, knockdown of CFI_m_25 results in a decrease in MAT2A mRNA after methionine, and therefore SAM, depletion. Additionally, knockdown of CFI_m_25 results in a reduction of intracellular SAM levels in a DI-dependent manner. Although the CFI_m_ complex has a widespread role in APA, it appears the regulation of SAM is separable from this function. We identified a CFI_m_25 binding sites in the 3′ UTR and one in the DI that are necessary for efficient splicing of MAT2A DI. Finally, we show that CFI_m_ induction of splicing requires the RS domains of CFI_m_68 or CFI_m_59. This leads to an updated model of splicing for the SAM-synthetase MAT2A, in which METTL16 serves as an upstream SAM sensor that mediates splicing of MAT2A through the CFI_m_ complex.

## RESULTS

### A CRISPR screen identifies CFI_m_25 as a candidate MAT2A splicing regulator

To investigate the mechanism of METTL16-mediated splicing of MAT2A, we designed a SAM-responsive GFP reporter cell line. Our reporter construct consists of GFP, a β-globin intron with flanking exonic regions, the MAT2A DI with flanking exons, and full-length MAT2A 3′UTR driven by a CMV promoter (Fig 1A). We integrated the reporter into the AAVS1 safe harbor locus on chromosome 19 of HCT116 cells and isolated single cell clones (Golden et al., 2017; Manjunath et al., 2019; Oceguera-Yanez et al., 2016). In SAM-replete conditions, the MAT2A intron should be inefficiently spliced and produce little GFP protein, while SAM depletion should lead to efficient splicing and robust GFP signal. To validate our reporter line, we depleted intracellular SAM using CRISPR/Cas9 with an sgRNA targeting the endogenous MAT2A. As expected, MAT2A knockout increased GFP signal (Fig 1B). Moreover, depletion of methionine, and therefore SAM, robustly increased GFP as monitored by flow cytometry (Fig 1C) and western blot (Fig S1A, “Reporter”). GFP-MAT2A mRNA accumulates upon shift to 11 μM methionine mirroring the endogenous MAT2A mRNA (Fig 1D). Similarly, knockdown of METTL16 abrogates the cell’s ability to induce production of the endogenous MAT2A or reporter mRNA after methionine depletion (Fig 1E). Together, these observations demonstrate that the production of our reporter mRNA mirrors that of the endogenous MAT2A.

Despite its usefulness in our studies, we noted that the reporter mRNA from our clonal line unexpectedly splices the β-globin 5′ splice site to the MAT2A 3′splice acceptor (Fig S1B-C). Other isolated clonal lines that spliced as predicted produced less GFP protein upon methionine depletion, the majority of which was accumulated as a putative degradation product (Fig S1A and S1D). Since the reporter splicing creates a distinct C-terminal extension on GFP, we reasoned that the GFP C-terminal extension produced from β-globin and MAT2A exons 8 and 9 destabilizes the GFP reporter protein (Fig S1B-C). Indeed, treating cells with the proteasome inhibitor MG132 increased protein levels produced in lines that had the predicted splicing pattern, but GFP protein was unaffected in our reporter line (Fig S1D). Thus, the alternative splicing of the reporter line results in a more stable protein product and therefore more useful reporter. The other difference between our reporter RNA and endogenous MAT2A RNA is that the polyadenylated DI isoform does not accumulate. The reason for this is unclear, but it may be related to relative stability and or 3′ processing efficiency of the unspliced RNA. Nevertheless, our reporter robustly responds to intracellular SAM levels in a METTL16-dependent fashion, mimicking the endogenous MAT2A. Therefore, we used this reporter cell line in a screen for factors required to induce MAT2A expression upon methionine depletion.

To identify factors necessary for induction of MAT2A splicing, we performed a CRISPR knockout screen. We reasoned that if we knock out genes essential for induction of MAT2A splicing (e.g. METTL16), cells will have reduced ability to induce GFP expression upon methionine depletion. We transduced our reporter line with the puromycin-resistant lentiviral Brunello library, which contains four sgRNAs per protein-coding gene of the entire genome (Doench et al., 2016). Two days post-transduction, we added puromycin, selected over 6 days, then replaced media with methionine-free media. Eighteen hours later, we sorted for the lowest ∼1% of GFP intensity cells (Fig 1F).

We identified seventy-two genes passing a 1% FDR cutoff from three independent biological replicates analyzed by MAGeCK (Fig 1G; Table S1)(Li et al., 2014). However, nearly all 72 genes were found on chromosome 19, where the AAVS1 safe harbor locus resides. The simplest explanation for this overrepresentation is that we selected for recombination events that reduced GFP signal by removing the reporter. After exclusion of genes on chromosome 19, three hits remained. The top candidate was METTL16 which supports the efficacy of the screen. The other hits were MED9 and NUDT21 (CPSF5). MED9 is a member of the mediator complex, a coactivator of RNA pol II (Soutourina, 2018). NUDT21 encodes the CFI_m_25 protein. Given its association with RS domain-containing proteins CFI_m_68 and CFI_m_59, we decided to investigate potential functions for CFI_m_25 in the regulated splicing of MAT2A.

### CFI_m_25 is required for induction of MAT2A mRNA and for maintenance of intracellular SAM levels

To validate the CRISPR screen, we depleted CFI_m_25 with two independent siRNAs in our reporter cell line. Upon knockdown of CFI_m_25, we observed a decreased ability to induce GFP and MAT2A mRNAs after methionine depletion (Fig 2A). To confirm this change is not cell line specific, we knocked down CFI_m_25 in 293A-TOA cells and analyzed MAT2A expression by northern blot analysis (Fig 2B). CFI_m_25 depletion increased intron detention under methionine-free conditions, phenocopying METTL16 depletion (Fig 2B, red). Interestingly, there was also a slight but significant increase in intron detention in methionine-replete media (Fig 2B, blue), further supporting a role for CFI_m_25 in the regulation of MAT2A intron detention. CFI_m_25 knockdown did not affect METTL16 expression (Fig 2C), so CFI_m_25 does not regulate MAT2A expression indirectly by manipulation of METTL16 abundance. Finally, we assessed SAM levels in cells depleted of CFI_m_25 and found that, like METTL16, depletion of CFI_m_25 decreased intracellular SAM levels in methionine-replete conditions (Fig 2D). Together, these data support the hypothesis that that CFI_m_25 is necessary for production of SAM by regulating MAT2A splicing.

**Figure 2.**
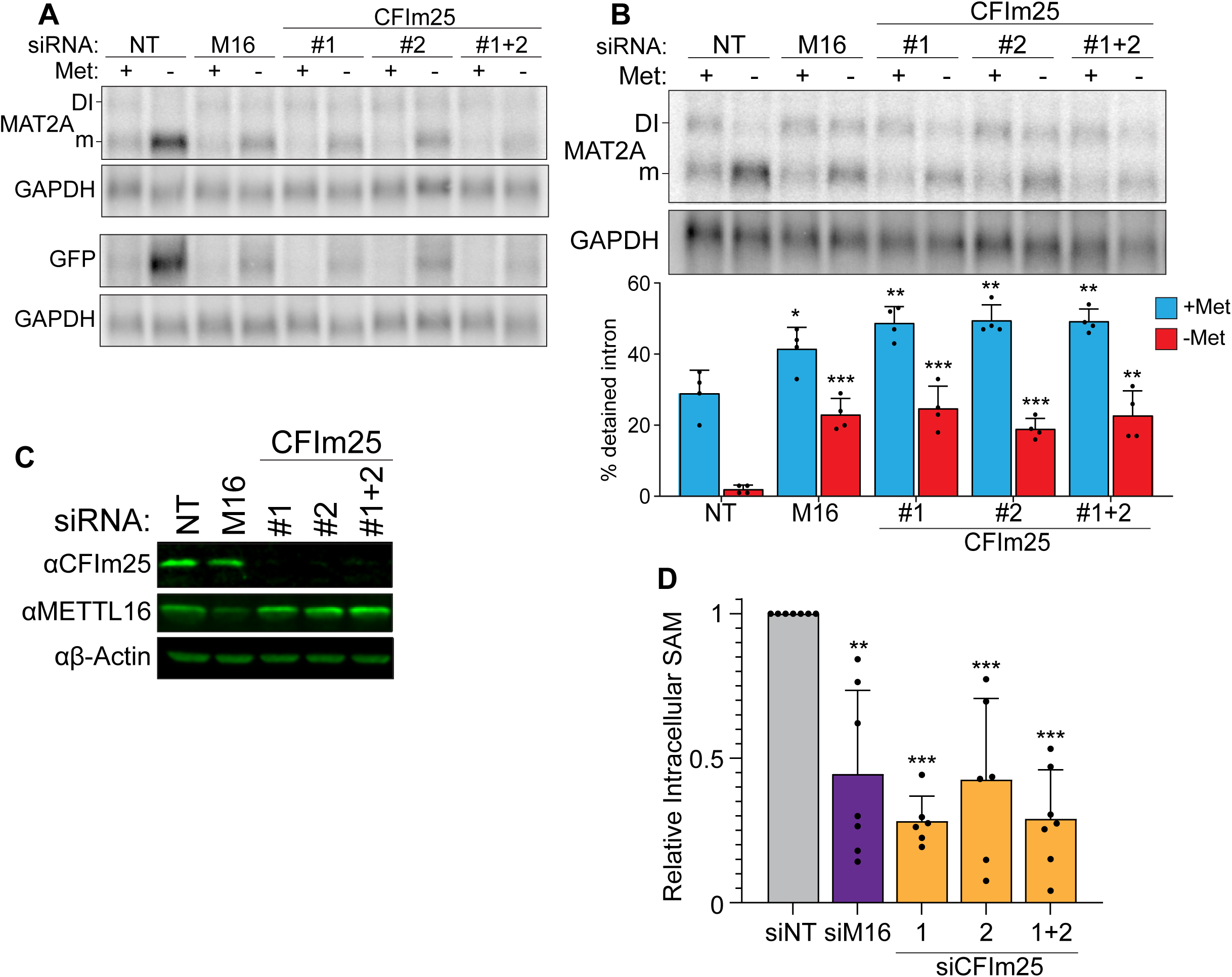
**CFI_m_25 regulates MAT2A Splicing and Activity** A. Northern analysis of MAT2A and GFP expression after knockdown with non-targeting (NT), METTL16 (M16), or two CFI_m_25 siRNAs individually or together in the reporter cell line. Cells were conditioned with methionine-rich or -free media for four hours before harvesting. B. Northern analysis of MAT2A expression after knockdown with non-targeting (NT), METTL16 (M16), or CFI_m_25 siRNAs individually or together in 293A-TOA cells. Cells were conditioned in methionine-rich or -free media for four hours before harvest. A representative blot is shown with quantification below. Asterisks denote statistical significance to the matched non-targeting sample; n=4. C. Western analysis of CFI_m_25 and METTL16 after the indicated knockdown in 293A-TOA cells. Actin serves as a loading control. n ≥ 3. D. Intracellular SAM levels relative to non-targeting control after METTL16 or CFI_m_25 knockdown in 293A-TOA cells. Statistical analysis compared all knockdowns to non-targeting control. Data are mean ± SD; n ≥ 6. Unless otherwise noted, data represented as mean ± SD and analyzed by a two-tailed, unpaired student’s t-test compared to matched control. Significance is annotated as not significant (ns), *p ≤ 0.05, **p ≤ 0.01, or ***p ≤ 0.001.

Knockdown of CFI_m_25 results in global changes in poly(A) site usage. Indeed, a minor isoform of MAT2A using a weak, proximal poly(A) site in the 3′UTR has previously been detected (Routh et al., 2017). Therefore, we tested whether CFI_m_25 regulates MAT2A by APA in our cells. A shorter isoform was not evident in our total RNA northern blots, so we poly(A) selected RNA for higher sensitivity. Two low-abundance bands of higher mobility were observed in 293A-TOA cells, either or both of which could be an APA product (Fig S2A). RNase H mapping results were consistent with one isoform being the previously reported APA isoform, but the longer isoform was of unclear origin (Fig S2B). Importantly, neither was responsive to methionine levels nor depletion of METTL16 or CFI_m_25 (Fig S2A). APA patterns can be cell-type specific (see below), but these data support the conclusion that the observed changes in MAT2A expression and SAM levels in 293A-TOA cells (Fig 2B-2D) are not due to CFI_m_- mediated changes in MAT2A poly(A) site usage.

### CFI_m_25 regulation of MAT2A requires the detained intron

Depletion of CFI_m_25 alters poly(A) site selection on many transcripts, so the observed changes in MAT2A and SAM levels may result from processes unrelated to MAT2A intron detention. Conversely, it is formally possible that some of the changes in APA are secondary to the drops in SAM levels observed upon CFI_m_25 depletion (Fig 2D). To test the importance of the DI, we used CRISPR to produce a clonal HCT116-derived cell line with the MAT2A DI deleted. Although we used a donor plasmid to promote homologous repair (HR) upon cleavage of two sites flanking the DI (Fig 3A), the resulting clonal line (116-ΔDI) is heterozygotic. One allele contains the expected precisely deleted DI from HR while the second allele deletes the DI via non-homologous end joining (NHEJ). The latter allele replaces the C-terminal 39 amino acids with 18 amino acids produced from a frameshift. As expected, no MAT2A-DI isoform was detected by northern blot, but methionine depletion leads to an increase of MAT2A mRNA presumably due to hp2-6 mediated stabilization of the mRNA (Fig 3B). Additionally, CFI_m_25 depletion had no effect on the methionine responsiveness of MAT2A mRNA in 116-ΔDI cells, consistent with a function for CFI_m_25 in regulation of the MAT2A DI (Fig S3A). We next compared intracellular SAM levels in 116-ΔDI to the parental line. Similar to 293A-TOA cells (Fig 2D), SAM levels decreased upon METTL16 or CFI_m_25 depletion in HCT116 cells (Fig 3C). In marked contrast, these treatments did not decrease SAM levels in 116-ΔDI cells. This observation strongly supports the conclusion that CFI_m_25 regulates SAM by regulation of the MAT2A DI.

**Figure 3.**
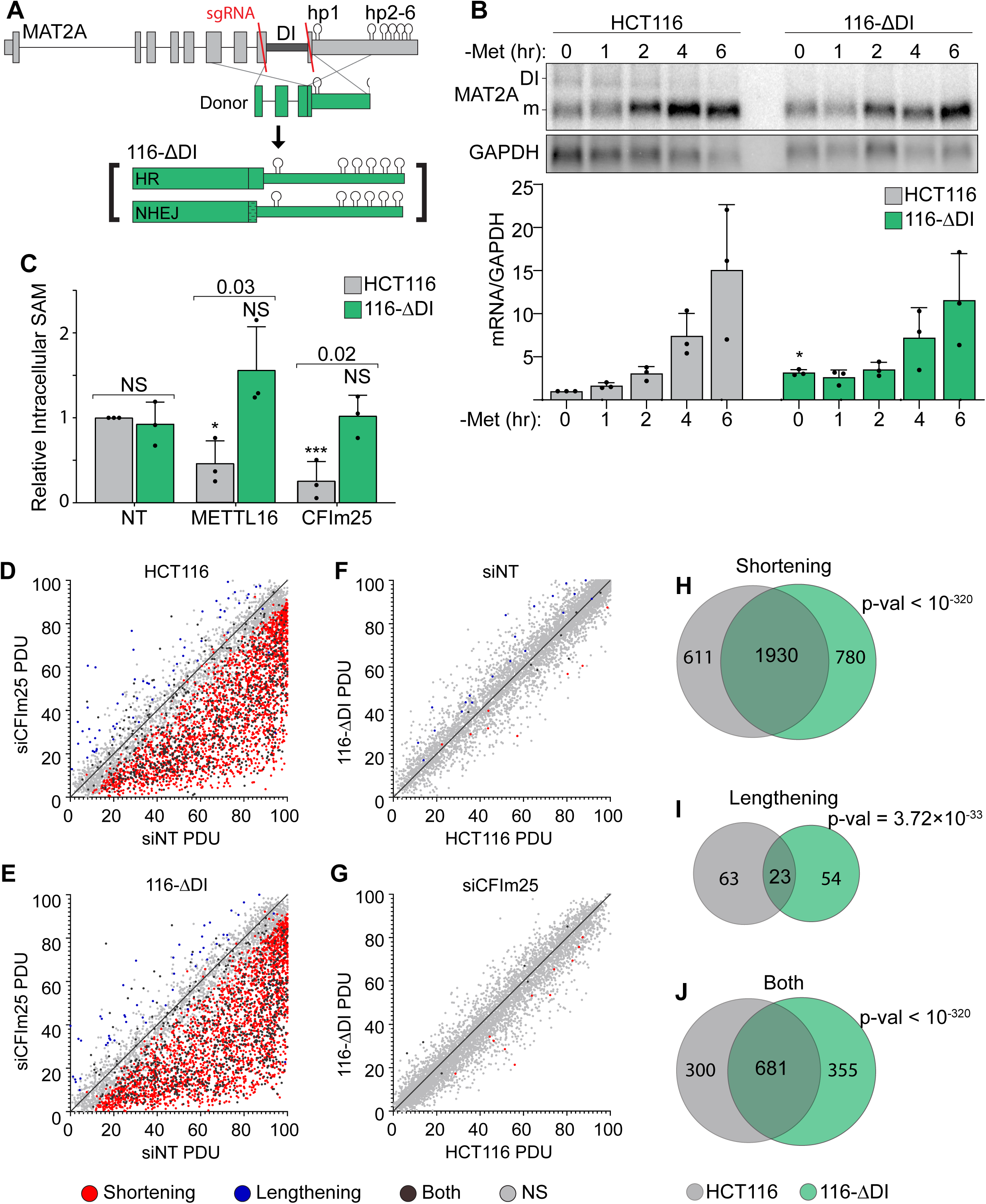
**CFI_m_25’s roles in APA and SAM Regulation are Separable** A. Schematic of the HCT116 ΔDI (116-ΔDI) cell line. Endogenous MAT2A gene was cleaved with Cas9 and two sgRNAs (red lines). One allele was repaired with an HR donor plasmid and the other was created by NHEJ. B. Northern analysis of MAT2A expression in the HCT116 parental and 116-ΔDI cell lines after the stated methionine depletion times. MAT2A mRNA levels were normalized to GAPDH and values are relative to HCT116 at 0 hr. Data are mean ± SD; n = 3. Statistics compare 116-ΔDI to HCT116 parental at the same time point. C. Intracellular SAM levels of HCT116 and 116-Δ DI cell lines after a 4-day knockdown with the indicated siRNAs. All values are relative to HCT116 parental non-targeting control. Two statistical comparisons are shown. Significance relative to the matched cell type non-targeting control is annotated with asterisks or NS. The p-values listed above the bars compare the two cell types within each knock-down condition. D-E. APA patterns in HCT116 parental and 116-ΔDI upon CFI_m_25 depletion. siCFI_m_25 vs siNT for HCT116 parental (D) and 116-ΔDI (E) plotted by percent distal site usage (PDU). Each dot represents a exon with multiple poly(A) clusters, with statistically significant shortening (red), lengthening (blue), or both (dark gray) APA events. Light gray dots are not statistically changed between samples (NS). F-G. Same as D-E, except HCT116 parental and 116-ΔDI cell lines were compared under siNon-targeting (F) and siCFI_m_25 (G) conditions. H-J. Venn diagrams comparing genes with shortening (H), Lengthening (I), or complex APA (Both, J) under CFI_m_25 depletion for HCT116 (gray) and 116-ΔDI (green) cell lines. P-values calculated using SuperExactTest (Wang et al., 2015).

In principle, some of the phenotypes associated with CFI_m_25 depletion may be due to its regulation of SAM. Since CFI_m_25 depletion does not alter SAM levels in 116-ΔDI cells (Fig 3C), we can use these cells to decouple CFI_m_25’s role in SAM metabolism and APA. To assess APA, we performed Poly(A)-ClickSeq (PAC-seq) that uses click chemistry techniques to sequence the 3′ ends of polyadenylated transcripts (Elrod et al., 2019; Routh, 2019; Routh et al., 2017). We detected over 2500 poly(A) site changes due to CFI_m_25 depletion (Table S2). Consistent with previous reports that depletion of CFI_m_25 favors the use of proximal poly(A) sites, the overwhelming majority of changes were reductions in the percent distal usage (PDU) of poly(A) sites (Fig 3D). Importantly, nearly identical levels of shortening were observed in 116-ΔDI (Fig 3E). In fact, when we compared the parental and 116-ΔDI lines under either knockdown DI (Fig 3H-J). Because the APA condition, only 45 and 20 genes were differentially polyadenylated for the siNT and siCFI_m_25, respectively (Fig 3F-G). Additionally, many exons with significant changes in APA were consistent between HCT116 and 116-ΔDI (Fig 3H-J). Because the APA patterns were largely similar even though the SAM levels differ upon CFI_m_25 knockdown (Fig 3C), these global analyses support our conclusion that CFI_m_25’s role in SAM regulation and APA are separable. They also strengthen the conclusion that CFI_m_25 regulates MAT2A mRNA abundance in a MAT2A DI-dependent fashion.

Consistent with our analysis in 293A-TOA cells (Fig S2), the PAC-seq analysis showed little use of the weak proximal MAT2A poly(A) site (Fig S3B and Table S2). Surprisingly, there was a statistically significant increase in use of this site in 116-Δ cells upon CFI_m_25 knockdown, although the distal poly(A) site is still the predominant isoform (Fig S3B and Table S2). In addition, there was a novel, albeit low abundance, poly(A) site in intron 7 in the 116-ΔDI cells only. We confirmed the existence of these RNAs by northern blot (Fig S3C-E). In principle, this could contribute to the restoration of MAT2A mRNA and protein because use of this site excludes the regulatory hp2-6. However, the low levels of each of these isoforms and their lack of induction in 293A-TOA cells (Fig S2A and S3) suggest they are not major contributors to MAT2A regulation in our experimental conditions. Thus, while these data may point to a more complex regulatory interface between splicing and MAT2A APA, they nonetheless support the conclusion that CFI_m_25 affects MAT2A processing independent of its well-described roles in APA.

### Two cis-acting CFI_m_25 binding sites regulate splicing of the MAT2A DI

To further test the role of CFI_m_25 in MAT2A splicing, we examined publicly available CLIP-seq data to determine if CFI_m_ interacts with the MAT2A 3′UTR (Martin et al., 2012). Because the preferred binding site for CFI_m_25 is UGUA (Brown and Gilmartin, 2003), we initially focused on this motif. All three components of the CFI_m_ complex cross-linked to varying degrees to the eleven UGUA motifs downstream of hp1 in the MAT2A 3′UTR (Fig 4A, ds-UGUA, blue hexagons). To determine if any of these consensus CFI_m_25 binding sites affect the splicing of MAT2A, we employed a reporter construct consisting of MAT2A exon 8, the DI, and exon 9 to the end of the 3′UTR fused downstream of β-globin coding sequence with an efficiently spliced β-globin intron (Fig 4B)(Pendleton et al., 2017). Using this construct, we first tested a mutation in the last motif (LM, UGUA to UG<u>C</u>A), which had the clearest binding in the CLIP-seq data overlapping a UGUA motif (Fig 4A). Because this mutation had no effect on intron detention (Fig 4C-D, dark blue), we then mutated ten out of eleven sites found in the 3′UTR to <u>C</u>GUA, U<u>C</u>UA, or UG<u>C</u>A (Fig 4C, 10/11). We alternated downstream mutations to prevent bias by introducing the same motif multiple times. The tenth site in the 3′UTR was not mutated due to overlap with the conserved METTL16 binding site in hp3 (Fig 4A, S4A)(Doxtader et al., 2018; Parker et al., 2011; Pendleton et al., 2017). The 10/11 construct had similar splicing efficiency as wild type, suggesting the downstream UGUA motifs are not involved in MAT2A splicing (Fig 4C-4D, light blue).

**Figure 4.**
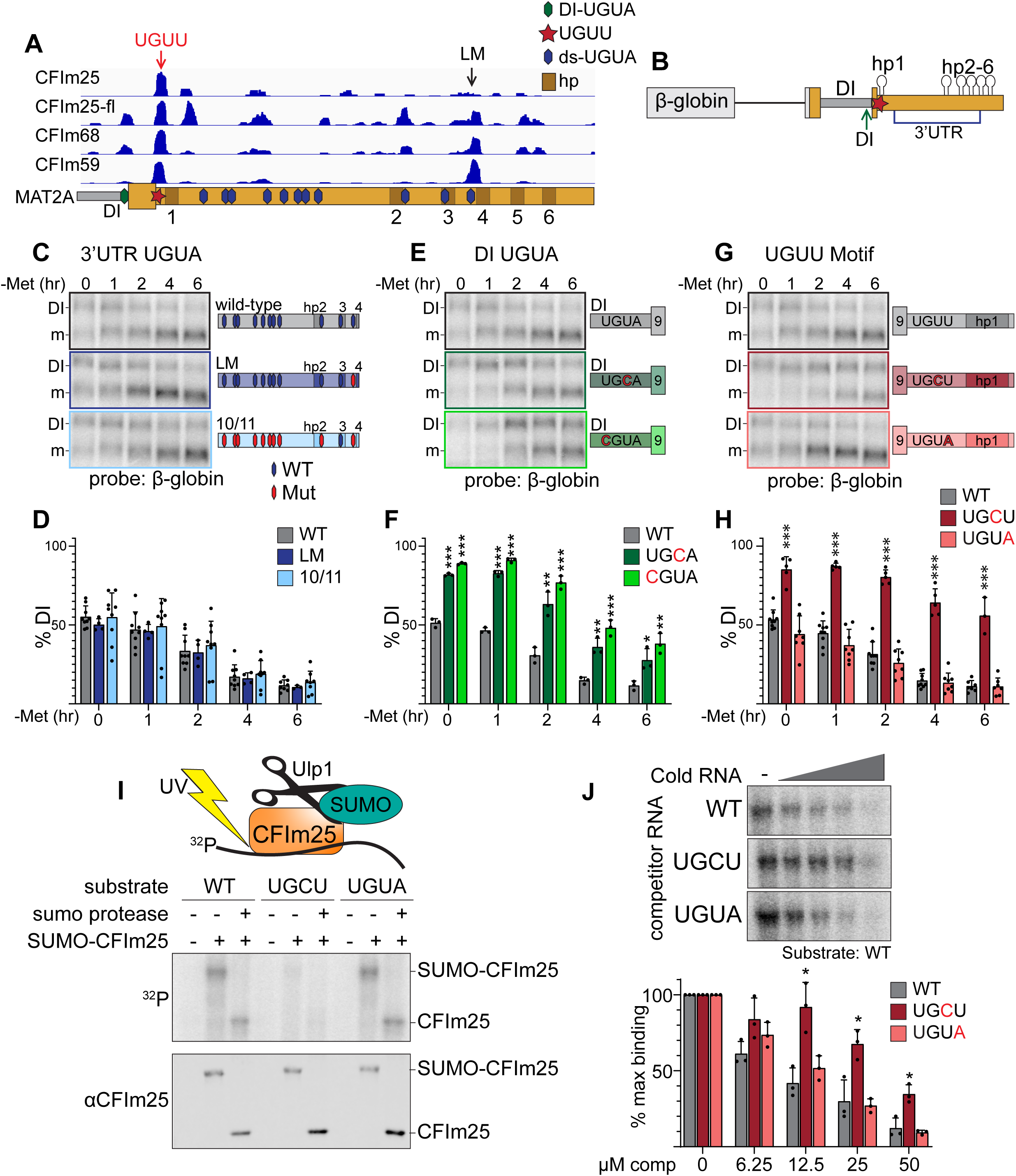
**A CFI_m_25 binding site in the DI and a noncanonical UGUU motif in the MAT2A 3′UTR are necessary for efficient MAT2A splicing.** A. IGV browser screen shot of CLIP-seq data of endogenous CFI_m_25, CFI_m_68, CFI_m_59, and flag-tagged CFI_m_25 (CFI_m_25-fl) to the MAT2A 3′UTR (Martin et al., 2012). Hexagons overlaid on the MAT2A schematic represent UGUA sites (blue, 3′UTR; green, DI UGUA). The UGUU-centered peak is denoted by a red star and red arrow. The last UGUA motif (LM) in the 3′UTR is denoted by a black arrow. MAT2A hairpins are denoted by brown boxes labeled 1-6. B. Schematic of the β-globin gene MAT2A reporter. The reporter consists of a β-globin gene excluding the first intron but maintaining intron 2 fused to MAT2A exon 8 through the end of the 3′UTR. Point mutations were categorized by potential binding site location. Blue bracket denoted 3′UTR includes the eleven downstream UGUA motifs (C and D). Green arrow denoted DI, the detained intron UGUA (E and F). Red star, the CLIP peak containing a UGUU (G and H). C and D. Representative northern blot and quantification of β-globin expression using reporters mutating UGUA motifs in the 3′UTR. In the schematics, hexagons represent UGUA elements, blue hexagons are wild-type, red are mutant. Gray, wild-type reporter (wt). Dark blue, mutation of the LM only (LM). Light blue, 10 of 11 dsUGUA motifs mutated (10/11). n ≥ 3. Representative northern blot data in Figure 4C, E, and G are from the same blot at the same exposure. Wild-type samples were run only once on that gel but the image is duplicated in the figure for easy comparison within each subgroup. E and F. Representative northern blot and quantification of β-globin expression using reporters mutating UGUA motif in the MAT2A DI. Gray, wild-type reporter (wt). Green, DI mutants UG<u>C</u>U (dark green) and <u>C</u>GUA (light green). n ≥ 3. G and H. Representative northern blot and quantification of β-globin expression using reporters mutating the UGUU motif immediately upstream of hp1. The UGUU was mutated to UG<u>C</u>U or to the canonical CFI_m_ binding motif (UGU<u>A</u>). Gray, wild-type reporter(wt). Dark red, UG<u>C</u>U. Pink, UGU<u>A</u>. n ≥ 3. I. Representative label transfer assay. SUMO-CFI_m_25 was incubated with radiolabeled 21-nt wild-type substrate centered on the UGUU within the natural sequence; two point mutants, UG<u>C</u>U and UGU<u>A</u> were also tested. In vitro binding was performed +/- SUMO-CFI_m_25 and +/- Upl1 SUMO protease as indicated. Top panel is label transfer, bottom is a western blot showing SUMO-CFI_m_25 loading in each lane. J. Competition label transfer assay. SUMO-CFI_m_25 was incubated with radiolabeled wild-type substrate (UGUU) plus increasing concentrations of cold wild-type or mutant substrate (UG<u>C</u>U, UGU<u>A</u>). Concentrations of competitor increase from left to right (0, 6.25, 12.5, 25, 50 μM). Gray, WT competitor. Red, UG<u>C</u>U competitor. Pink, UGU<u>A</u> competitor. n=3.

An evolutionarily conserved UGUA motif is found in the DI 9-12 nt upstream of the 3′splice site (Fig 4A, S4B). Due to its proximity to the polypyrimidine tract in the 3′splice site, we tested two independent point mutations (UG<u>C</u>A, <u>C</u>GUA) that maintain pyrimidine content but disrupt the CFI_m_25 binding motif. We found that both mutations significantly abrogated MAT2A splicing both at basal levels and upon methionine depletion (Fig 4E-F). Thus, intronic assembly of the CFI_m_ complex may contribute to MAT2A splicing.

The strongest and most consistent CLIP-seq peak among all three CFI_m_ factors lies immediately upstream of hp1 and downstream of the stop codon (Fig 4A). There is no UGUA element, but the peak centers on a UGUU motif that is evolutionarily conserved in vertebrates, except in *Danio rerio* where it is a UGUA, the canonical CFI_m_25 binding site (Fig S4C). We mutated the UGUU to UG<u>C</u>U which decreased splicing efficiency comparably to that of the DI point mutations (Fig 4G-H, dark red). We observed no changes in splicing efficiency when the UGUU motif was replaced with the canonical UGUA sequence (Fig 4G-H, pink). Thus, this non-consensus binding site for CFI_m_ appears to contribute to the splicing of the MAT2A DI.

Consistent with the CLIP-seq and our functional data, structural and biochemical analysis of CFI_m_ interactions with RNA suggest some flexibility in the fourth position of the UGUA motif (Yang et al., 2011; Yang et al., 2010). To further validate that CFI_m_ binds to the UGUU site, we conducted label transfer assays using SUMO-tagged recombinant CFI_m_25. Radiolabeled 21-mer RNA substrates containing the UGUU or its variants were incubated with SUMO-CFI_m_25, cross-linked, then analyzed by SDS-PAGE. We found that CFI_m_25 crosslinked the substrates containing either the canonical UGU<u>A</u> or UGU<u>U</u> motifs more efficiently than to the UG<u>C</u>U substrate (Fig 4I). The addition of SUMO protease Upl1 to remove the SUMO tag increased band motility to further confirm that the band represents crosslinked CFI_m_25-RNA complexes. To further test the relative binding of these RNAs to CFI_m_25, we performed a competition assay in which radiolabeled wild-type UGUU substrate was competed with cold UGUU, UG<u>C</u>U, and UGU<u>A</u> RNAs. Cold UG<u>C</u>U RNA was a significantly less efficient competitor than UGUU or UGU<u>A</u> RNAs (Fig 4J). Thus, the simplest explanation for the reduced splicing in the UGCU reporters (Fig 4G-H) is that CFI_m_25 binds this motif less efficiently than the UGUU or UGU<u>A</u> motifs (Fig 4I-J). Additionally, UGUU and UGU<u>A</u> appear to bind CFI_m_25 and function within MAT2A to comparable levels (Fig 4G-I), suggesting the UGUU motif is not necessarily a weak CFI_m_25 binding site despite its deviance from the well-studied UGU<u>A</u> motifs. Together, the CLIP-seq (Martin et al., 2012), reporter assays, and *in vitro* binding all suggest that CFI_m_25 binds to a noncanonical UGUU motif in MAT2A’s 3′UTR to regulate intron detention.

### The CFI_m_ complex is required for induction of MAT2A splicing

Both CFI_m_25 cofactors CFI_m_68 and CFI_m_59 contain RS domains, so we hypothesized that CFI_m_68 and/or CFI_m_59 promote the splicing of MAT2A. Depletion of CFI_m_59 with two different siRNAs individually or in combination had no significant effects on MAT2A intron detention, while depletion of CFI_m_68 had modest effect for only one siRNA (#1; Fig 5A, 5B). In contrast, co-depletion of CFI_m_68 and CFI_m_59 increased intron detention upon methionine depletion compared to the matched individual knockdowns (Fig 5A-B). Only modest effects of CFI_m_68 and CFI_m_59 depletion were observed in methionine-replete conditions (Fig S5A). CPSF73 (CPSF3), a CFI_m_- independent component of the cleavage and polyadenylation complex, served as an additional control and showed no change in intron detention (Chan et al., 2011). In addition, MAT2A intron detention is nearly identical after exposing cells to media with different methionine concentrations upon CFI_m_68 and CFI_m_59 co-depletion, METTL16, or CFI_m_25 depletion (Fig 5C-D). These results suggest possible functional redundancy between CFI_m_68 and CFI_m_59 in the splicing of MAT2A, consistent with the lack of identification of either gene in our CRISPR screen (Fig 1H). However, due to co-dependent protein stability among the members of the CFI_m_ complex, variable levels of co-depletion of factors occur (Fig S5B)(Chu et al., 2019). Therefore, these data alone do not conclusively show that MAT2A splicing requires CFI_m_68 and CFI_m_59.

**Figure 5.**
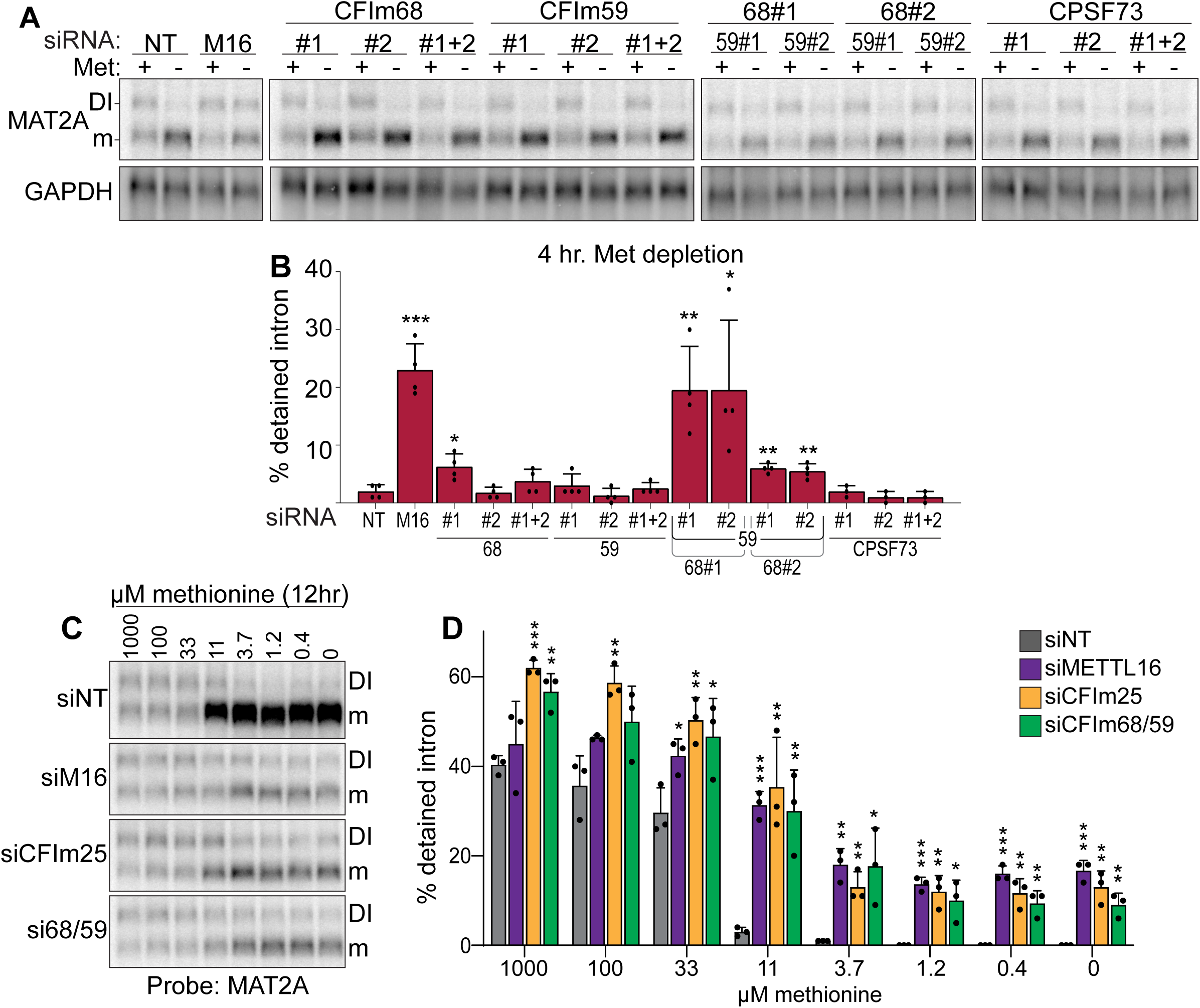
**Co-depletion of CFI_m_68 and CFI_m_59 reduces induction of splicing of the MAT2A DI.** A-B. Representative northern blot and quantification of MAT2A intron detention after CFI_m_68 and CFI_m_59 (co-)depletion. Two independent siRNAs for each factor were used (labeled #1 and #2). 293A-TOA cells were conditioned with methionine-rich or methionine-free media for 4 hrs prior to harvesting. n ≥ 3. C-D. Representative northern blot and quantification of MAT2A expression after methionine titration after depletion of the indicated factors. 293A-TOA cells were conditioned with media containing the methionine concentration specified for 12 hrs prior to harvesting. Non-targeting control, siNT, gray. METTL16, siM16, purple. siCFI_m_25, orange. siCFI_m_68 and siCFI_m_59 co-depletion, green. n = 3.

To further test a direct role for CFI_m_68 and CFI_m_59 in MAT2A splicing induction, we performed a series of tethering assays. We employed a MAT2A β-globin reporter with two bacteriophage MS2 coat protein binding sites immediately downstream of hp1. Hp1 in this construct contains an A4G mutation that abrogates METTL16 binding and induction of splicing (Fig 6A)(Pendleton et al., 2017). We co-expressed MS2-coat protein fusions to determine their effects on MAT2A splicing when artificially tethered to the reporter RNA in cells. First, we tethered wild-type CFI_m_25 (wt) under methionine-rich conditions and observed induced splicing of the reporter to levels comparable with METTL16 tethering (Fig 6B). Importantly, both the MS2-NLS-fl alone and the reporter lacking MS2-binding sites generated low levels of spliced product. We next tested two CFI_m_25 mutants that bind neither CFI_m_68 nor CFI_m_59. The first construct consists of two point-mutations in the nudix hydrolase domain (m1, Y158A/Y160A) and the second contains a single point mutation near the C-terminus (m2, L218R)(Fig 6D)(Yang et al., 2011; Zhu et al., 2018). We confirmed that CFI_m_25 binding to CFI_m_68 and CFI_m_59 was abrogated in both mutants by coimmunoprecipitation (Fig 6C). Upon tethering of these CFI_m_ mutants, we observed a significant loss in splicing induction suggesting that a functional CFI_m_ complex is responsible for splicing (Fig 6B). Importantly, the CFI_m_25 mutants expressed to comparable levels to that of the wild-type construct (Fig 6C input). In contrast, tethering of an RNA-binding mutant (m3, R63S) of CFI_m_25 maintained splicing activity, as expected since RNA binding is driven by MS2 tethering and it maintains binding to CFI_m_68 and CFI_m_59 (Fig 6c)(Yang et al., 2011; Yang et al., 2010).

**Figure 6.**
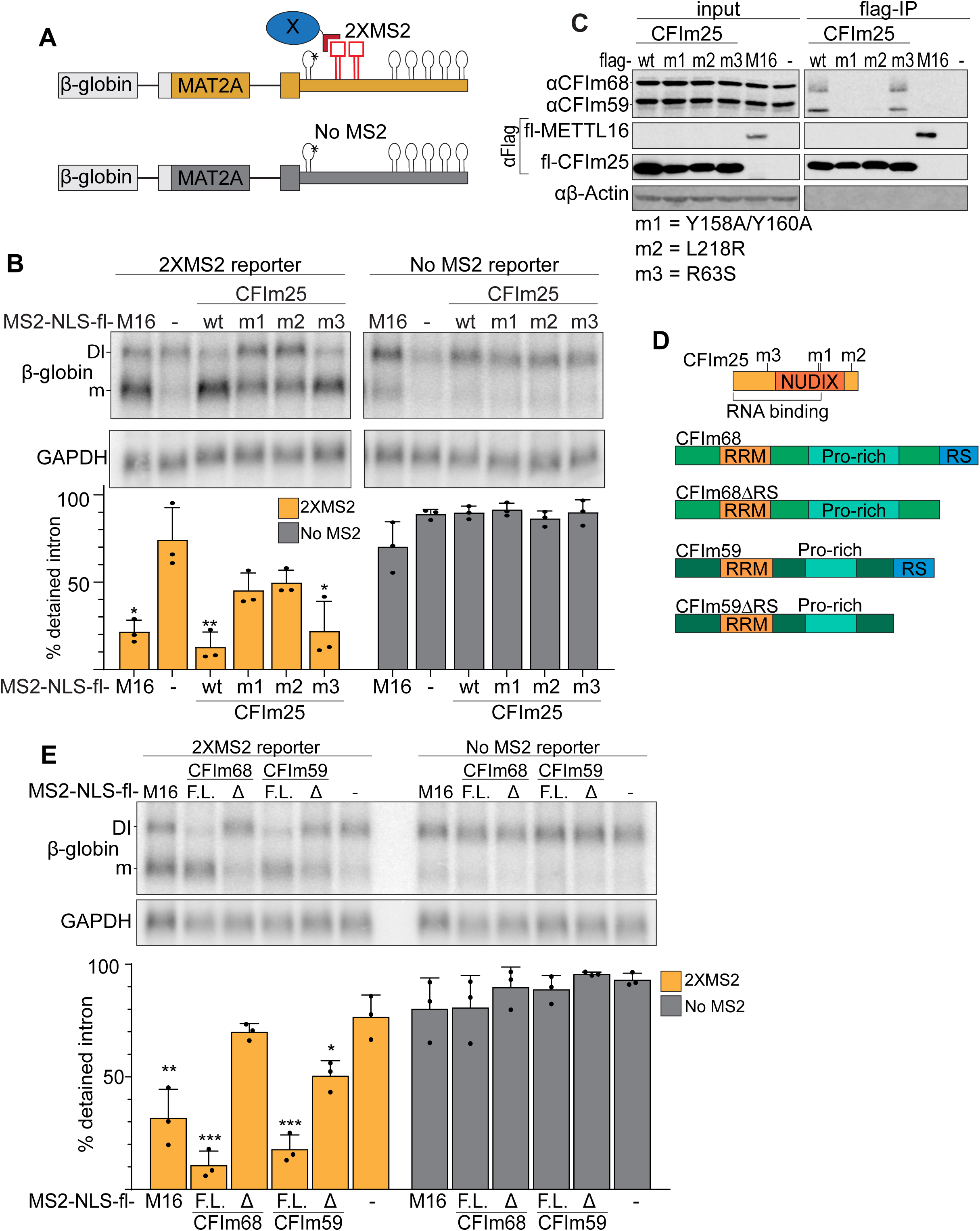
**Binding of the CFI_m_ complex is sufficient to promote MAT2A splicing.** A. Diagram of the tethering assay. The 2XMS2 β-globin reporter consists of MAT2A exon 8, the detained intron, exon 9, and the full-length 3′UTR with two bacteriophage MS2-coat protein binding sites inserted 3′ of A4G mutant hp1 (asterisk). A matched reporter lacking the MS2 site (“No MS2”) is used to measure background. All MS2 fusion proteins have an N-terminal MS2 coat protein, an SV40 nuclear localization signal, and Flag tag (MS2-NLS-fl). The MS2-NLS-fl alone is expressed as negative control. Diagram not to scale. B. Northern analysis of β-globin reporter RNA after tethering CFI_m_25 variants. MS2-NLS-fl fusions to METTL16 (M16), CFI_m_25 variants (wt, m1, m2, m3), or MS2-NLS-fl alone (-) and were expressed with the indicated reporters. Statistical analysis is relative to matched MS2-vector control. Orange, 2XMS2 reporter. Gray, No MS2 reporter. n = 3. C. Coimmunoprecipitation of CFI_m_68 and CFI_m_59 with flag-CFI_m_25. Flag-tagged wild-type CFI_m_25 (wt) or mutants (m1, m2, m3), flag-tagged METTL16 (M16), or flag-vector (-) were expressed in HEK293 cells before immunoprecipitation with anti-flag beads. The immunoprecipitates were then probed for endogenous CFI_m_68 and CFI_m_59, flag, or β-actin. Input is 10% of the lysate volume applied to flag beads. n=3. D. Diagrams of CFI_m_25, CFI_m_68, CFI_m_68Δ RS, CFI_m_59, and CFI_m_59Δ RS proteins; diagrams to scale. E. Northern analysis of β-globin reporter RNA after tethering CFI_m_68 or CFI_m_59. MS2-NLS-flag tagged METTL16, CFI_m_68, CFI_m_68ΔRS, CFI_m_59, CFI_m_59ΔRS or vector (-) were co-transfected with the 2XMS2 β-globin reporter or no MS2 site control. Statistical analysis is relative to matched MS2-vector control. F.L., full length; Δ, ΔRSdomain. n = 3.

CFI_m_25 has no known splicing domains, however, both CFI_m_68 and CFI_m_59 contain N-terminal RS domains. Therefore, we reasoned that the RS domains of CFI_m_68 and CFI_m_59 may be responsible for the splicing of MAT2A. If so, tethering full-length CFI_m_68/CFI_m_59 will induce splicing of MAT2A, while the tethering of CFI_m_68/CFI_m_59 lacking the RS domains (ΔRS) will not (Fig 6D). As predicted, CFI_m_68 and CFI_m_59 tethering was sufficient to induce splicing, while the Δ proteins were unable to affect intron detention (Fig 6E). Lack of splicing was not due to inability of the constructs to express, as the RS domain deletion expressed to comparable levels of their full-length counterparts (Fig S6A). Together, these observations suggest that the CFI_m_ complex is responsible for the splicing of the MAT2A DI.

### CFI_m_ is downstream of METTL16 in the MAT2A splicing induction mechanism

Our data suggest the model for CFI_m_ complex-mediated induction of MAT2A splicing shown in Figure 7A. CFI_m_ is recruited to a UGUA in the DI and UGUU in MAT2A 3′UTR. Upon CFI_m_ binding, CFI_m_68 and CFI_m_59 serve as the downstream effectors that promote efficient splicing of the DI through their RS domains. If CFI_m_ is indeed the effector complex, its function in splicing lies downstream the SAM sensor METTL16. To test the proposed hierarchy of factors in the splicing of the MAT2A DI, we combined our tethering system with siRNA knockdown of individual factors. In some cases, the expression of the MS2 transgenes is reduced upon knockdown of these essential factors (Fig S7). Importantly, in each of these cases, our data show that the lower levels remain sufficient to potently drive reporter splicing (see Fig S7 legend). Therefore, the MS2 fusions are overexpressed at levels such that slight changes in their expression does not confound our results. If CFI_m_ is downstream of METTL16 as suggested, tethering METTL16 will no longer induce splicing upon CFI_m_25 depletion. Indeed, the tethering of METTL16 fails to induce splicing after CFI_m_25 depletion compared to the non-target and no MS2 controls (Fig 7B). Conversely, tethering of CFI_m_25, so long as it is in complex with CFI_m_68 or CFI_m_59, should induce splicing even in METTL16-depleted cells. As expected, tethering of CFI_m_25 after METTL16 depletion results in splicing induction comparable to the non-targeting control (Fig 7B). This observation is also consistent with the ability of CFI_m_ complex members to induce splicing in the hp1 A4G mutant, which no longer binds METTL16 (Fig 6). Together, these observations support the idea that METTL16 acts as the upstream SAM sensor while CFI_m_ directly mediates the splicing of MAT2A.

**Figure 7.**
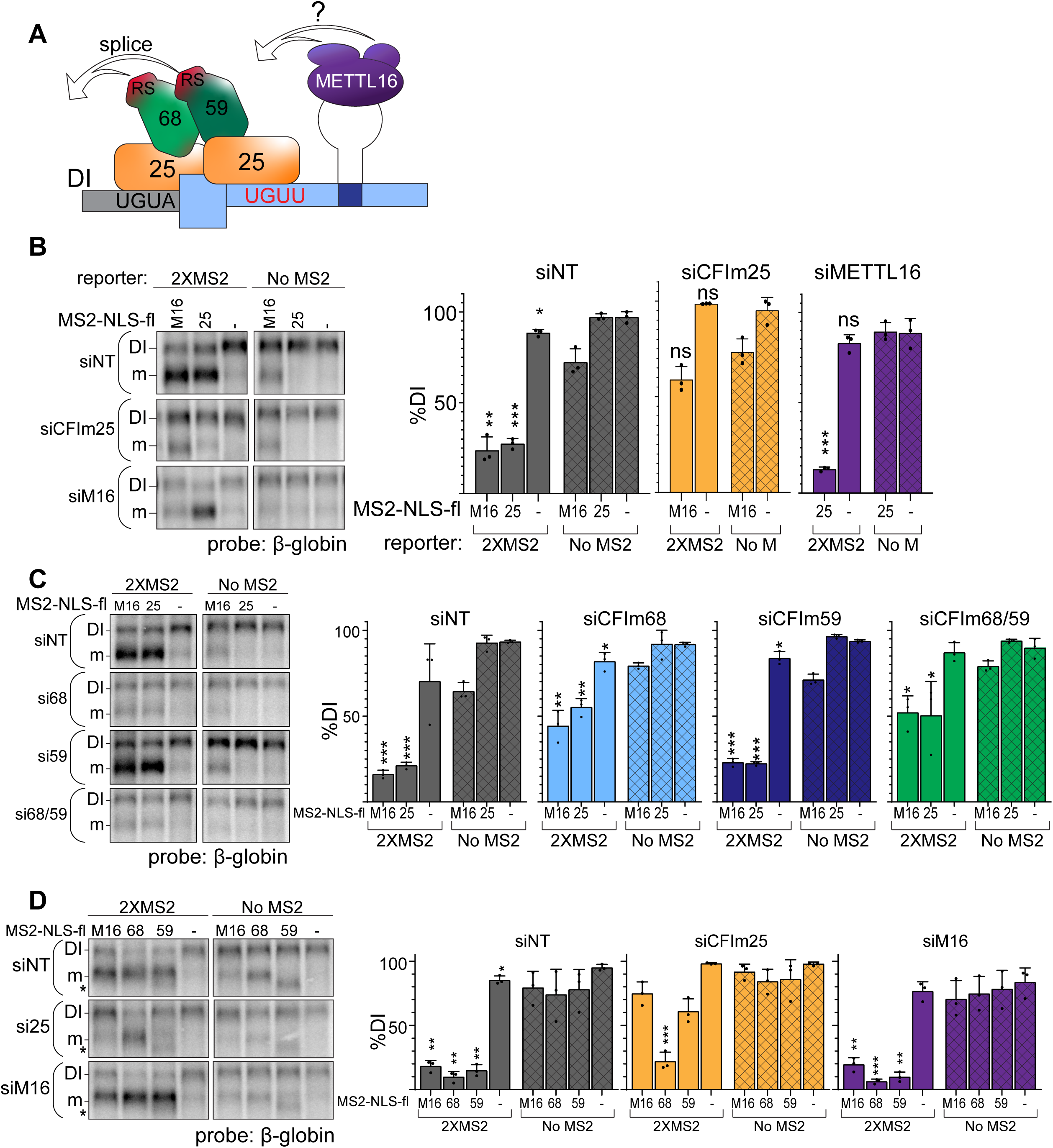
**CFI_m_ is the downstream splicing effector of METTL16** A. Model for CFI_m_-induced MAT2A splicing. CFI_m_ complex binds to a non-canonical UGUU motif in the MAT2A 3′UTR and UGUA in the detained intron. This allows for MAT2A splicing by proximity of the RS domains in CFI_m_68 or CFI_m_59. How this is integrated with METTL16 binding remains unknown (see Discussion). B. Northern analysis of β-globin reporter RNA after knockdown of the indicated factor and tethering of MS2-NLS-fl tagged METTL16 (M16), CFI_m_25 RNA binding mutant (m3, 25), or MS2-NLS-fl vector (-). *Left*, representative northern blot. *Right*, quantification by percent detained intron for non-targeting siRNA (siNT, gray), or siRNAs targeting CFI_m_25 (si25, orange) or METTL16 (siM16, purple). n = 3. C. Same as (B). Non-targeting siRNA (siNT, gray), siCFI_m_68 (si68, light blue), siCFI_m_59 (si59, dark blue), and siCFI_m_68/59 co-depletion (si68/59, green). n = 3. D. Same as (B) except tethering of MS2-NLS-fl-METTL16 (M16), -CFI_m_68 (68), - CFI_m_59 (59), or MS2-NLS-fl vector (-) with the MS2-reporter constructs in cells depleted of METTL16 or CFI_m_25. Overexpression of CFI_m_59 caused a band of unknown identity to appear (asterisk). The band was cell-type specific: it did not appear in HEK293 cells (Fig 6E), but it does in 293A-TOA cells used here. Non-targeting siRNA (siNT, gray), siCFI_m_25 (si25, orange), and siMETTL16 (siM16, purple). n = 3. Statistical analysis in B-D compares 2XMS2 reporter to matched, no-MS2 reporter.

Our model proposes that CFI_m_25 recruits CFI_m_68/CFI_m_59 to the RNA and CFI_m_68 or CFI_m_59 interchangeably function in the splicing of MAT2A. If so, tethering METTL16 or CFI_m_25 will no longer induce splicing if the downstream effectors CFI_m_68 and CFI_m_59 are not present. Consistent with this prediction, CFI_m_68/CFI_m_59 co-depletion reduces splicing induction upon METTL16 or CFI_m_25 tethering (Fig 7C, green). Surprisingly, depletion of CFI_m_68 alone had similar effects as CFI_m_68/CFI_m_59 co-depletion (Fig 7C, blue), while CFI_m_59 depletion was comparable to the non-targeting control (Fig 7C, dark blue). We likewise observed that tethering CFI_m_68 induces splicing in CFI_m_25-depleted cells, but CFI_m_59 did not (Fig 7D). These data demonstrate that tethering CFI_m_68 is sufficient to induce splicing of MAT2A in the absence of METTL16 or other CFI_m_ complex family members. The function of CFI_m_59 in MAT2A splicing is less clear, but it may contribute to splicing redundantly with CFI_m_68 or have other functions in MAT2A splicing or CFI_m_ complex assembly (see Discussion).

## Discussion

We propose that the CFI_m_ complex, a major APA factor, is a key regulator of SAM metabolism. Next to METTL16, CFI_m_25 was the top candidate in a global unbiased screen for regulators of MAT2A splicing. Knockdown and tethering experiments independently support the conclusion that components of the CFI_m_ complex drive splicing of the MAT2A DI through the RS domains of CFI_m_68 and/or CFI_m_59. Moreover, mutation of CFI_m_25 binding sites in the MAT2A DI and 3′UTR both abrogate splicing. Taken together, these data support the model shown in Figure 7A. We conclude that the CFI_m_ complex is the downstream effector that promotes MAT2A splicing regulated by the SAM sensor METTL16.

This model leads to an important remaining question: how is METTL16 binding to hp1 connected to CFI_m_-mediated MAT2A DI splicing? The simplest model is that METTL16 recruits CFI_m_ to the RNA, but we were unsuccessful in coimmunoprecipitating METTL16 and CFI_m_25. Additionally, our unpublished yeast two-hybrid experiments and co-immunoprecipitation mass spectrometry experiments were consistent with a published report that found no clear METTL16 binding proteins (Ignatova et al., 2019). It seems unlikely that METTL16 functions without other binding partners, but if METTL16 and CFI_m_25 directly interact, the interaction is likely transient. Alternatively, METTL16 may mediate CFI_m_25 binding to MAT2A through an RNA conformational change that exposes the binding sites. This conformational change hypothesis is supported by Aoyama et al.’s discovery that the VCR domain of METTL16 binds RNA (Aoyama et al., 2020). Since the VCR domains alone can drive splicing when tethered to a reporter (Pendleton et al., 2017), the VCR domain could alter RNA structure surrounding hp1 to promote CFI_m_25 binding (Aoyama et al., 2020). Alternatively, other RNA, proteins, or posttranslational modifications may promote the transient formation of the METTL16:CFI_m_ complex on the MAT2A transcript. Such proteins would likely function redundantly with other factors because they were not identified in the CRISPR screen. In any case, future experiments are required to distinguish among these and other models that explain the interface between METTL16 and CFI_m_ in MAT2A splicing.

We have determined that the CFI_m_ complex is necessary for the splicing of MAT2A DI, but the composition of the complex that promotes splicing is unclear. In our knockdown experiments CFI_m_68 and CFI_m_59 appear to be redundant for splicing (Fig 5) and neither appeared as CRISPR screen hits (Fig 1, Table S1). Further supporting functional redundancy, tethering of either CFI_m_68 or CFI_m_59 resulted in comparable splicing induction, as long as an RS domain is present (Fig 6E). Despite CFI_m_68 and CFI_m_59 belonging to the same complex and sharing significant sequence and structural similarity, CFI_m_68 and CFI_m_59 have non-redundant functions in cells (Deng et al., 2019; Li, 2010a, b; Tresaugues, 2010; Yang et al., 2011). Knockdown of CFI_m_68 leads to a global shift to proximal poly(A) site usage, similarly to CFI_m_25 knockdown, while CFI_m_59 has no effect (Gruber et al., 2012; Zhu et al., 2018). Additionally, CFI_m_68 and CFI_m_59 have distinct interaction partners (Martin et al., 2012; Martin et al., 2010; Millevoi et al., 2006). Although CFI_m_68 and CFI_m_59 initially appear to be functionally similar for the splicing of MAT2A, some of our data minimize the role of CFI_m_59. When tethered, CFI_m_25 is incapable of promoting splicing in the absence of CFI_m_68, while CFI_m_59 knockdown alone has no effect (Fig 7C). Additionally, tethering of CFI_m_68 enables splicing independent of CFI_m_25, while CFI_m_59 requires CFI_m_25 (Fig 7D). One explanation is rooted in the three possible compositions of the CFI_m_ complex. The CFI_m_25 dimer can form a tetramer with two CFI_m_59 proteins, two CFI_m_68 proteins, or with one of each. Thus, tethering of CFI_m_59 may lead to recruitment of the CFI_m_ complex that includes CFI_m_68. If so, then tethering of a CFI_m_25:CFI_m_68:CFI_m_59 containing complex would be capable of promoting splicing but would require CFI_m_25 to recruit CFI_m_68. More testing needs to be done to unravel the overlapping and distinct roles of CFI_m_68 and CFI_m_59 in the splicing of MAT2A.

CFI_m_ binding sites in the MAT2A DI and immediately upstream of hp1 are necessary for MAT2A splicing. Mutation of either the DI UGUA or 3′UTR UGUU results in abrogation of splicing to comparable levels, suggesting that both sites contribute equally to the splicing of MAT2A (Fig 4E-H). The two sites could function as independent CFI_m_ binding sites. Alternatively, since CFI_m_25 exists as dimer within the complex, it is possible that the sites are bound simultaneously with the intervening RNA looped out. In fact, previous data suggests looping may be important for the function of CFI_m_25 (Yang et al., 2011; Yang et al., 2010). Two UGUA motifs in the PAPOLA mRNA 3′UTR simultaneously bind to the CFI_m_25 dimer to promote more efficient binding. The two sites must be at least 9 nt apart to support looping, but the upper limit of distance between UGUA motifs has yet to be tested. The UGUA in the MAT2A DI and the UGUU upstream of hp1 are 114 nt apart allowing ample separation for RNA looping. It has been proposed that RNA looping mediated by the CFI_m_ complex regulates poly(A) site selection by occluding or presenting poly(A) sites (Yang et al., 2011). However, to our knowledge, this has yet to be formally demonstrated in cells. Given that one of the sites is in the DI near the 3′splice site (Fig S4C), it is tempting to speculate that CFI_m_-dependent RNA looping exposes the 3′splice site to increase its accessibility to the spliceosome.

The CFI_m_ complex is a major regulator of poly(A) site selection (Brumbaugh et al., 2018; Gruber et al., 2012; Li et al., 2015; Martin et al., 2012; Masamha et al., 2014; Zhu et al., 2018). Our data show that the CFI_m_ complex’s role in MAT2A splicing and SAM metabolism appear to be independent of poly(A) site selection (Fig 3). The shortening of 3′UTRs upon CFI_m_ knockdown has been linked to important biological phenotypes including cancer cell proliferation and cell differentiation (Brumbaugh et al., 2018; Chu et al., 2019; Jafari Najaf Abadi et al., 2019; Masamha et al., 2014; Tamaddon et al., 2020). For example, knockdown of either CFI_m_25 or CFI_m_68 increases transcription-factor-induced reprogramming of mouse embryonic fibroblasts (MEFs) into induced pluripotent stem cells (iPSC)(Brumbaugh et al., 2018; Ji and Tian, 2009). This effect is attributed to global shifts from distal poly(A) site usage to proximal, which is characteristic of less differentiated and more proliferative cells (Ji et al., 2009; Mayr and Bartel, 2009; Sandberg et al., 2008; Shepard et al., 2011). Our work shows that SAM levels likely drop in MEFs upon CFI_m_ knockdown, so it is possible that SAM reduction contributes to the increased dedifferentiation potential by priming cells for a transition into iPSC (Shiraki et al., 2014). Similarly, it is plausible that SAM levels contribute to the growth potential of cancer cells, as cancer cells often have reduced methylation potential and/or intracellular SAM levels (Gao et al., 2019; Murray et al., 2019). Therefore, CFI_m_ knockdown may augment cancer growth by reducing SAM levels in addition to the well-defined 3′UTR shortening mechanism (Jafari Najaf Abadi et al., 2019). Thus, it will be interesting to decouple CFI_m_25’s role in poly(A) site selection from that in maintenance of intracellular SAM to see how each contributes to these important biological phenomena.

## Acknowledgements

We thank Drs. Feng Zhang, David Root, John Doench, Didier Trono, and Kathryn Pendleton for plasmids. We also would like to thank Drs. Joshua Mendell and John Schoggins and their lab members for advice in designing the reporter cell line and conducting CRISPR screens. CRISPR screens were sorted with the help of the Moody Foundation Flow Cytometry Facility in the Children’s Research Institute at UTSW. This research was supported by the Welch foundation I-1915-20170325 (to N.K.C), the National Institutes of Health NIGMS: R01 GM127311 to N.K.C.; R01 GM127311-S1 to J.N.F.; T32 GM007062 to A.M.S.; R35 GM136370 to B.P.T..

## Author Contributions

Conceptualization: N.K.C and A.M.S; Methodology: A.M.S, J.N.F., O.V.H., K.L., A.K., C.X., B.P.T., N.K.C.; Investigation: A.M.S., J.N.F., K.L., O.V.H., and N.K.C.; Data Curation: A.K., C.X.; Writing – Original Draft: A.M.S. and N.K.C.; Writing – Review & Editing: A.M.S., J.N.F., K.L., A.K. B.P.T., N.K.C.; Supervision: C.X., B.P.T., and N.K.C.; Funding Acquisition: C.X., B.P.T., and N.K.C.

## STAR Methods

### Resource Availability

#### Lead Contact

Further information and reagent requests may be directed to Nicholas K. Conrad.

#### Materials Availability

Plasmids generated in this study are available upon request of the lead contact, Nicholas K. Conrad.

#### Data and Code Availability

Raw and unedited CRISPR screen data is available upon request. MAGeCK analysis of CRISPR screen data is found in Table S1.

Raw and unedited Poly(A)-ClickSeq data is deposited on GEO (GSE158591). Analysis of Poly(A)-ClickSeq is found in Table S2.

### Experimental Models and Subject Details

#### Cell culture

HEK293 and HEK293T cells were maintained in DMEM (Sigma, Cat#D5796) with penicillin-streptomycin, 2 mM L-glutamine, and 10% fetal bovine serum (FBS, Sigma, Cat#F0926) and grown at 37°C in 5% CO_2_. The same media was used with a maintenance concentration of Plasmocin™ (InvivoGen, Cat#ant-mpt, 1:10,000) for all HCT116-based cell lines and 50 μg/ml hygromycin for reporter cell lines. 293A-TOA cells were cultured similarly to HEK293 cells, but with Tet-free FBS (Atlanta Biologicals, Cat#S10350) and 100 μg/ml G418. Care was taken to ensure cells were passaged in methionine-rich media to keep MAT2A DI/mRNA ratios consistent between experiments. Methionine-free media DMEM (Thermo Fisher, Cat#21013024) was supplemented with 0.4 mM L-cysteine and 1 mM sodium pyruvate in addition to penicillin-streptomycin, L-glutamine, and Tet-free FBS.

#### Generation of GFP-reporter line

HCT116 cells were co-transfected in a 6-well plate with 0.2 µg AAVS1 1L TALEN, 0.2µg AAVS1 1R TALEN, and 1.6 µg hAAVS1-GFP-β2-MAT-E8-3′. The next day cells were split to 10 cm plates, and hygromycin was added to 100 µg/ml. Cells were selected for a total of two weeks, initially in 100 µg/ml hygromycin and then in 250 µg/ml hygromycin over the second week. Fluorescence-activated cell sorting (FACS) was used to select clonal cell lines with low to mid GFP output in methionine-rich conditions. Clonal cell lines were selected based on high differential GFP expression between methionine-rich and methionine-starved conditions. The cell line that provided the most robust GFP expression after methionine depletion was chosen as the reporter line. As described (Fig S1), the robust differential results in part from an alternate splicing pattern that stabilizes the GFP protein.

#### Generation of 116-ΔDI line

Two 10 cm plates of HCT116 cells were each transfected with 3 µg of pX458-MAT2A-E9, 3 µg of pX459-MAT2A-E8, and 10 µg of pBS-ΔRI-Donor. Eight hrs later, fresh media was added that included 1 µg/ml puromycin and 1 µM of the NHEJ inhibitor SCR7 (Fisher Scientific; Chu et al., 2015; Maruyama et al. 2015). Approximately 48 hrs after transfection, puromycin-resistant transfected cells were subjected to FACS, and GFP-positive single cells were seeded on a 96-well plate. SCR7 was included for an additional 3-5 days. After clonal expansion, DNA was harvested, and MAT2A DI status was examined by PCR using primers NC1145 and NC2537. The sequences of all primers and DNA oligonucleotides are listed in Table S3. Only one clonal line contained DI deletions on both alleles. Subsequent sequencing demonstrated that one allele had the DI deleted by HR while the other allele had intron 8 deleted by NHEJ. The latter deleted an additional 20 nt (18 nt from exon 8 and 2 nt from exon 9) to create an out-of-frame junction between exons 8 and 9: CGA TCT CCG**/AT CTG GAT** (exon8/**exon9;** see also Fig 5A).

### Method Details

#### Methionine depletion

Cells were transferred into fresh DMEM media supplemented with an additional 200 μM methionine the day before depletion. To deplete methionine, cells were washed twice with phosphate buffered saline (PBS) supplemented with calcium chloride and magnesium chloride (Sigma) before replacing with growth media containing or lacking methionine as required.

#### Transfection

Cells were transfected using TransIT-293 (Mirus, Cat#MIR 2706) according to the manufacturer’s protocol. For co-immunoprecipitations, 600 μl Opti-MEM™ (Thermo Fisher, Cat#31985-070) and 36 μl TransIT-293 were incubated at room temperature (RT) for 5 min before addition to 10 μg of aliquoted DNA. The mixture was incubated 15 min before adding dropwise to a 10 cm dish of cells. For typical 12-well transfections, 2 μ l Opti-MEM, and 800 ng plasmid was added per well using the above protocol. MS2-tethering experiments used 30 ng of reporter, 200 ng of MS2-NLS-flag tagged construct, and 570 ng of pcDNA3 per well of a 12-well plate. Expression of MS2-NLS-flag protein was analyzed after transfecting 400 ng of MS2-NLS-flag construct and 400 ng pcDNA per well. In some cases, HCT116 cells were transfected using FuGENE® HD Transfection Reagent (Promega, Cat#E2311) according to the manufacturer’s protocol.

#### RNA extraction and purification

RNA was harvested using TRI Reagent® (Molecular Research Center, Inc., Cat#TR 118) with minor variations to manufacturer’s protocol. For one well of a 12-well plate, 400 μ l chloroform was added to the TRI Reagent containing tube before shaking vigorously by hand until homogenization. The samples were centrifuged at 12,000 x g for 15 min at 4°C before transferring the aqueous phase to a fresh tube then mixing with an equal volume of chloroform. Care was taken not to disturb the interphase between the organic and aqueous phases. The chloroform/aqueous mixture was shaken vigorously by hand and centrifuged at 12,000 x g for 5 min at RT before transferring the aqueous phase once again to a fresh tube. The aqueous solution was mixed with an equal volume of isopropanol for storage at –20°C or precipitation by centrifugation at 16,000 x g 10 min RT with the addition of 15 μg GlycoBlue™ Coprecipitant (Thermo Fisher, Cat#AM9516).

#### siRNA knockdown

293A-TOA or HCT116 lines and their derivatives were transfected with 30 nM siRNA using RNAiMAX following the manufacturer’s protocol. Twenty-four hrs after transfection, confluent cells were split to allow for an additional three days of cell division and knockdown (total 96-hr knockdown). Degree of cell dilution when passaging at this stage was dependent on the toxicity associated with the specific knockdown. We changed the media 24 hrs before harvesting to ensure cells were maintained in a methionine-rich conditions. For samples using multiple siRNAs targeting the same gene, 15 nM of each siRNA was used for a total of 30nM. For knockdown prior to tethering (Fig 7), cells were transfected with the appropriate tethering constructs 72 hrs post knockdown after a media change to ensure cells were harvested in methionine-rich conditions. Cells were harvested 24 hrs post-transfection and 96 hrs post-knockdown.

#### Poly(A) selection

Sera-Mag Oligo(dT)-Coated Magnetic particles (GE Healthcare Lifesciences, Cat#38152103010150) were washed three times in 1XSSC (150 mM NaCl, 15 mM sodium citrate) with 0.1% SDS then resuspended in the same using the initial volume. Purified total RNA in water was heated at 65°C for 5 min before the addition of SSC and SDS to 1X and 0.1% respectively. Washed Sera-Mag Oligo(dT)-Coated Magnetic particles were added to the RNA (20 μl particles per 40 μg total RNA). Samples were nutated 20 min at RT then washed three times in 0.5X SSC/0.1% SDS. RNA was eluted in 100 μl water for 5 min at RT. The supernatant was combined with a second elution in 100 μl water for 5 min at 65°C. The combined eluants were further purified by phenol:chloroform:iso-amyl alcohol (PCA, 25:24:1) extraction, chloroform extraction, and ethanol precipitation.

#### RNase H Mapping

Poly(A) selected RNA purified from an initial 160 μg total RNA in H_2_O was divided equally into three tubes. Five µM of specific DNA oligonucleotide and 1 μM dT_20_ was added, then samples were diluted to 10 μ Lreaction volumes before incubation at 65°C for 5 min. After cooling on ice 3 min, the following was added to each tube to reach a final volume of 20 µl per reaction: RNase H buffer (20 mM Tris pH7.5, 100 mM KCl, 10 mM MgCl_2_ final concentration), 10 mM DTT, 0.75 U RNase H, and 16U RNasin Plus. Samples were digested for 1 hr at 37°C before quenching with 180 μ l G50 buffer (20 mM Tris pH7, 300 mM sodium acetate, 2 mM EDTA, 0.25% SDS), then purified using PCA and chloroform extractions followed by ethanol precipitation.

#### Northern blotting

Northern blots were performed using standard techniques and probed with radiolabeled RNA transcripts (Ruiz et al., 2019). RNA probes were produced using a digested plasmid or PCR products containing a T7 or SP6 RNA polymerase promoter. Primer and plasmids are listed in Table S3. Either 3-5 μg total RNA or polyadenylated enriched RNA produced from 40 μg total RNA was loaded per lane.

#### Coimmunoprecipitation

Plasmids expressing flag-tagged proteins were transfected into 10 cm dishes of HEK293 cells. After washing twice, cells were scraped in PBS. Cells were pelleted at 1000 x g 4°C for 3 min. PBS was removed and cells were lysed in RSB100T (50 mM Tris pH 7.5, 100 mM NaCl, 2.5 mM MgCl_2_, 1 mM CaCl_2_, 1% TritonX100) with 1 mM PMSF and 1X Protease Inhibitor Cocktail Set V (Millipore, Cat#539137). Samples were nutated at RT with 20U RQ1 RNase-free DNase (Promega, Cat#M6101) and RNase A (10 μg/ml) for 15 min before clarifying twice by centrifugation at 4°C 21,000 x g for 10 min. Lysate was bound to pre-washed ANTI-FLAG® M2 Affinity Gel (Sigma, Cat#A2220) by nutating at 4°C for 2 hrs before washing five times with RSB100T. Protein was eluted by vortexing for 30 min at 4°C in RSB100T supplemented with 0.4 mg/ml 3X FLAG® peptide (Sigma, Cat#F4799), 1 mM PMSF and 1X Protease Inhibitor Cocktail Set V. Samples were analyzed by standard western blotting protocols.

#### SAM metabolite extraction

Protocol was adapted from Dettmer et al. (2011) and Tu et al. (2007). Cells were treated with siRNAs as indicated and maintained in methionine-rich conditions in 10 cm plates. Ninety-six hrs after knockdown, cells were washed three times with ice-cold PBS with calcium chloride and magnesium chloride before the addition of 600 μl ice-cold 80% methanol while being snap frozen in liquid nitrogen. Samples were scraped on ice, transferred to Eppendorf tube after mixing by pipetting, then frozen in liquid nitrogen. Samples were thawed in a RT water bath while mixing by pipetting before clarification at 16,000 x g 4°C for 10 min. Methanol supernatants were stored at -80°C and cell pellets were washed 1X PBS, resuspended in 1X SDS PAGE loading buffer without loading dye (2% SDS, 62.5 mM Tris pH 6.8, 10% glycerol, 1% β-mercaptoethanol) then sonicated until homogenous. Relative protein concentration measured by nanodrop was used to estimated cell number between samples and methanol supernatant volumes were adjusted accordingly. Methanol supernatants were dried using a speed vacuum before resuspension in Solvent A (1% formic acid in water), centrifuged twice, then filtered through 0.2 μM PVDF to remove insoluble particles. Samples were analyzed via LC-MS/MS with a total run time of 20 min, flow rate of 0.5 ml/min, 0.1% formic acid in water as Solvent A, and 0.1% formic acid in methanol as Solvent B. Pure SAM was injected and analyzed alongside samples for each experiment. SAM was detected by multiple reaction monitoring (MRM) using the ion pair 339/250, quantified using the Analyst 1.6.1 Software package by calculating total peak area, then normalized to non-targeting control (Dettmer et al., 2011)(Tu et al., 2007).

#### Flow cytometry

HCT116 reporter cells were conditioned in methionine-rich or methionine-free media for 24 hrs before harvesting by trypsinization. After quenching the trypsin, cells were pelleted by centrifugation at 800 x g for 3 min at 4°C then washed with PBS. After washing, cells were resuspended in 5% formaldehyde in PBS and nutated 1 hr to overnight. Prior to analysis by flow cytometry, cells were pelleted at 800 x g for 3 min at 4°C then resuspended in PBS with 3% FBS. Samples were aliquoted into 96-well v-bottom dishes to be analyzed on a Stratedigm S1000. Flow cytometry data was analyzed by FloJo to compare relative GFP fluorescence.

#### CRISPR screen

The Human Brunello CRISPR knockout pooled library, a gift from David Root and John Doench (Addgene #73179), was prepared according to the BROAD institute pDNA library amplification protocol (BROAD Institute Amplification of pDNA libraries; Doench et al., 2016; Shalem et al., 2014). In brief, 400 ng of library was electroporated into ElectroMAX™ Stbl4™ Competent Cells (Thermo Fisher, Cat#11635018). After recovery in SOC Outgrowth Medium (New England Biolabs, Cat#B9020S), the sample was plated equally between four 500 cm^2^ bioassay plates containing LB agar with 100 μg/ml puromycin using a biospreader (Bacti Cell Spreader, VWR International). Cells were grown 18 hrs at 37°C before scraping with ice cold LB. DNA was prepared by dividing the total cell mass evenly between two Qiagen maxi prep columns, following the manufacturer’s protocol.

To produce lentivirus, twenty 15 cm tissue culture plates were coated with poly-D-lysine (100 ug/ml in milliQ water, filtered) for 5 min then washed twice with PBS before plating 293T cells. pMD2.G and psPAX2 were a gift from Didier Trono (Addgene plasmid #12259 and #12260). Brunello library, pMD2.G, and psPAX2 were co-transfected between the twenty plates equally, using a total of 300 μg, 120 μg, and 180 μg plasmid respectively. Prior to transfection, the media was exchanged for DMEM supplemented with 3% FBS instead of the normal 10%, with subsequent media used for the production of lentivirus likewise containing 3% FBS. Media was changed 6 hrs post transfection, then collected and pooled at 48 and 72 hrs post-transfection. HEPES pH 7.2 (20 mM) and 4 ug/ml polybrene was added to the lentivirus before filtration through a 0.45 μ filter to clear debris. The filtered virus was treated with Benzonase® Nuclease (Sigma, Cat#E1014) to digest residual plasmid (50 U/ml benzonase, 50 mM Tris pH 8.0, 1 mM MgCl_2_, 0.1 mg/ml BSA) for 30 min at 37°C with gentle agitation. Aliquoted lentivirus was snap frozen in liquid nitrogen then stored at -80°C until further use.

Lentivirus titer was determined by transduction of HCT116 reporter cells plated at 4X10^5^ cells per well of a 6-well plate. Cells were split evenly into media ±1 ug/ml puromycin 48 hrs post-transduction. Five days after transduction, cells were harvested and analyzed via CellTiter-Glo® Luminescent Cell Viability Assay (Promega, Cat#G7570). To estimate the viral titer, we compared cell counts in selected vs unselected conditions for each lentivirus treatment condition.

For the genome-wide screen, a titer that infected ∼20% of plated cells was used. To obtain 100X coverage of the 76,441 gRNA in the Brunello library, sixteen 6-well plates with 4X10^5^ cells per well were infected. Two days post transduction, cells were split into media containing 1 μg/ml puromycin and Plasmocin™ (1:10,000). Media was changed every day and cells were split as needed, with 200 μM additional methionine added on day seven post-transduction. On day eight, cells were deprived of methionine 18 hrs before sorting by FACS. Care was taken to sort cells within an 18-20 hr window after methionine depletion to maintain consistency between replicates.

To prepare cells for FACS, cells were trypsinized then diluted in ice-cold PBS, pipetting gently yet thoroughly to ensure cell clumps were broken up. Cells were pelleted at 1,600 x g for 3 min at 4°C before resuspension in FACS buffer (50 mM HEPES pH 7.2, 0.5 mM EDTA, 2% Tet-free FBS, PBS). Resuspended cells were strained using a 100 μM nylon cell strainer (Falcon® 100 μM Cell Strainer, Cat#352360) and kept on ice before and during sorting. Cells with the lowest 1% of GFP signal were collected in FACS collection buffer (50% Tet-free FBS, 50 mM HEPES pH 7.2, PBS) with two rounds of enrichment using a BD Biosciences FACSAria Fusion Cell Sorter. The first round of enrichment of cells sorted at approximately 12,000 cells/sec. However, due to the droplet size, several mid to high GFP expressing cells appeared in the sample. The second round of enrichment removed the mid to high GFP expressing cells, only accepting cells within the initial gating set for the lowest 1% of GFP expressing cells relative to the unsorted population. For unsorted input to compared to the GFP-depleted sample, a minimum of 8 million cells were set aside for lysis without sorting. Sample coverage was calculated by the number of cells collected in the final sample divided by 1% of the number of gRNA found in the Brunello library (Equation 1). Cells were centrifuged at 1,600 x g for 3 min 4°C, resuspended in 500 μ l tissue lysis buffer (10 mM Tris pH 8, 100 mM NaCl, 11 mM EDTA, 200 ug/ml Proteinase K, 0.4% SDS) and lysed at 55°C 550 rpm overnight.

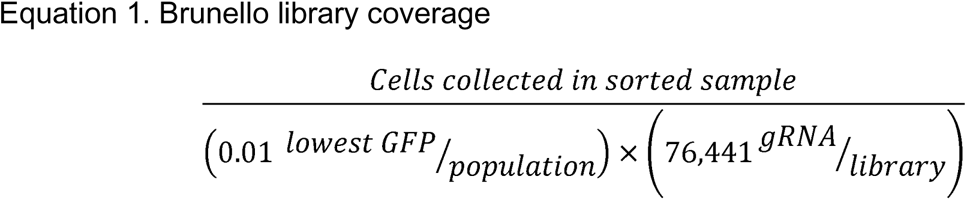

To isolate DNA, the lysates were cooled 3 min at RT before the addition of 5 μ 2 mg/ml RNase A. After briefly vortexing, samples were shaken at 37°C 550 rpm for 1 hr. An equal volume of PCA was added per tube before vortexing 20 sec and dividing equally between two prespun phase lock tubes (1.5 ml MaXtract High Density Tubes, Qiagen, Cat#129046). Samples were spun at 16,000 x g for 5 min RT before an additional 500 μl PCA was added to each phase lock tube, without transferring the aqueous layer. Tubes were inverting vigorously for ∼1 min to mix then centrifuged at 16,000 x g for 5 min RT. The aqueous phase was then transferred to a fresh, prespun phase lock tube with the addition of 500 μl chloroform. The tubes were once again inverted vigorously before centrifuging at 16,000 x g for 5 min. The aqueous phase was transferred to a fresh Eppendorf tube with the addition of 20 μ g GlycoBlue Coprecipitant. DNA was precipitated in ethanol at -80°C, pelleted at 16,000 x g RT for 10 min before being washed with 75% ethanol. DNA pellets were resuspended in sterile water at RT for 30 min. After adequate time for resuspension, the DNA was pipetted ∼20 times to shear the DNA.

Library amplification was completed following a two-step variation of the BROAD institute protocol (BROAD Institute PCR of sgRNAs for Illumina sequencing). The two-step variation of the BROAD institute protocol consists of an initial amplification step using primers flanking the P5 and P7 primers used for sequencing. The initial PCR product (PCR#1) is then diluted to be used as template for amplification Illumina P5/P7 flow cell primers (PCR#2). For the first PCR step, 6.6 μ g DNA was used as template per 100 μl reaction, with twenty reactions being set up per sample if possible. If the sample contained less than 13.2 μg DNA, the DNA was divided evenly between a minimum of two reactions. The DNA was amplified using with TaKaRa ExTaq and primers NC3196 and NC3197 flanking the P5 and P7 sites (1X ExTaq reaction buffer, 0.2 mM each dNTP, 0.5 μ g template DNA, water to 100 μl per reaction). PCR#1 consisted of 18 cycles following the ExTaq manufacturer’s protocol (95°C 1 min, [95°C 30 sec, 53°C 30 sec, 72°C 30 sec] x 18, 72°C 10 min, hold 4°C). For the second PCR amplification step, 5 μl of the pooled PCR#1 reaction was used as template for each 100 μl PCR#2 reaction. Four 100 μl reactions were made per sample using the reaction conditions above, except replacing the primers NC3196 and NC3197 with Illumina P5 stagger primer mix and a P7 barcode primer (Table S3). The P5 stagger primer consisted of all eight stagger primers evenly mixed. A unique P7 barcode primer was used for each sample. The PCR cycle number for PCR#2 was selected to match band intensity between samples when run on an agarose gel (8-12 cycles).

The 400 μl PCR#2 product per sample was pooled before purification by AMPure XP (Beckman Coulter, Cat#A63880) following the BROAD institute protocol (BROAD Institute PCR of sgRNAs for Illumina sequencing). Briefly, the pooled PCR volume was mixed with 0.5 AMPure XP bead volume then incubated 5 min RT. Beads were separated on a magnetic strip for 2 min then the supernatant transferred to a fresh tube to be mixed with 1.8 volumes of AMPure XP beads. After a 5 min RT incubation, beads were separated for 3 min on a magnetic strip before washing twice with 70% EtOH for 1 min each while remaining on the magnet. The beads were dried for 5 min before eluting with the starting volume of water (400 μl) for 2 min at RT. The beads were separated from the sample for 2 min on the magnetic strip, and the eluted DNA was transferred to a fresh tube. The purification protocol was repeated a second time, with the only change being a final elution volume of 50 μl water instead of 400µl.

Amplified library was analyzed by Qbit, TapeStation, and qPCR to assess library purity and concentration before sequencing. Three independent biological replicates were sequenced on an Illumina NextSeq 500 with read configuration as 75bp, single end. The fastq files were subjected to quality check using fastqc (version 0.11.2, http://www.bioinformatics.babraham.ac.uk/projects/fastqc) and fastq_screen (version 0.4.4, http://www.bioinformatics.babraham.ac.uk/projects/fastq_screen), and adapters trimmed using an in-house script. The Human Brunello CRISPR library sgRNA sequences were downloaded from Addgene (“http://www.addgene.org/pooled-library/)” www.addgene.org/pooled-library/). The trimmed fastq files were mapped to Brunello library sequence with mismatch option as 0 using MAGeCK (Li et al., 2014). Further, read counts for each sgRNA were generated and median normalization was performed to adjust for library sizes. Positively and negatively selected sgRNA and genes were identified using the default parameters of MAGeCK.

#### Poly(A)-ClickSeq

RNA was extracted from HCT116 or 116-ΔDI cells grown in methionine-rich conditions and treated with non-targeting or CFI_m_25 siRNA. RNA was treated with RQ1 RNase-free DNase with the addition of RNasin® Plus RNase Inhibitor (Promega, Cat#N2615)(DNAse RQ1 1X reaction: pH 8.0, 10 mM NaCl, 6 mM MgCl_2_, 10 mM CaCl_2_, 5U RQ1 DNase, 40U RNasinPlus) for 30 min at 37°C before PCA and chloroform extraction. Precipitated RNA was analyzed by TapeStation to verify that the RNA samples had a RIN of 8.5 or greater before submission.

Poly(A)-ClickSeq was performed by ClickSeq Technologies, Galveston, TX, using methods described elsewhere (Elrod et al., 2019; Routh, 2019; Routh et al., 2017). Briefly, samples were prepared by RT-PCR using an oligo-dT with a P7 adapter. The P5 primer was attached using click-chemistry before amplifying the RT product. The samples were sequenced on an Illumina NextSeq 550 using a Mid Output 130M, v2 flow cell. Only reads with >40nts of cDNA sequence and >10 A’s were retained for analysis to ensure mapping of the 3′UTR. The reads were quality filtered then aligned, with most localization to the 3′UTR of transcripts. Sites with >5 reads within 10nts were defined as a poly(A) cluster (Elrod et al., 2019; Routh et al., 2017). Samples were analyzed using Differential Poly(A)-Clustering (DPAC)(Routh, 2019).

#### Purification of recombinant SUMO-CFI_m_25

Rosetta (DE3) cells (EMD Millipore) were transformed with pE-SUMO-CFI_m_25 and selected in 30 μ g/ml kanamycin. Colonies were inoculated into a 2 mL starter culture and grown at 37°C overnight. The culture was diluted into 200 mL fresh LB media with antibiotics, grown to mid-log phase (O.D. ∼0.5), and IPTG was added to 1 mM. After 2 hr at 37°C, bacterial pellets were harvested by 10 min centrifugation at 4,000 x g and the pellets were resuspended in 1 mL lysis buffer (300 mM NaCl, 50 mM NaH_2_PO_4_, 0.5% Triton X-100, 0.5 mM PMSF, pH 8.0). Two milligrams lysozyme (Sigma) was added and the mix was incubated on ice for 30 min. Benzonase was then added to 0.25 U/mL and the mix was nutated for 30 min at RT and the lysate was cleared by centrifugation at 10,000 x g for 30 min at 4°C. The protein was purified in batch by incubation with 1 mL of Ni-NTA Agarose (QIAGEN, Cat#30210) for 1 hr at 4°C. The beads were washed over a column with ten volumes of wash buffer (300 mM NaCl, 50 mM NaH_2_PO_4_, 10 mM imidazole, 0.5% Triton X-100, 0.5 mM PMSF, pH 8.0) and proteins were collected in fractions of elution buffer (300 mM NaCl, 50 mM NaH_2_PO_4_, 250 mM imidazole, pH 8.0). Eluted fractions were pooled and dialyzed twice at 4°C in 2L of Buffer D (20 mM Hepes pH 7.9, 20% glycerol, 50 mM KCl, 0.2 mM EDTA, 0.5 mM DTT) using a Slide-A-Lyzer Dialysis Cassette (10,000 kD cutoff; Fisher). The first dialysis step was for 2 hr, buffer was replaced, and the second dialysis step occurred overnight. The samples were concentrated ∼2-fold using Amicon Ultra 0.5 centrifugal filter units to a final concentration 0.8 mg/mL. Aliquots of the protein were stored at -80°C.

#### Label transfer assays

The RNAs used as substrates and competitors were synthesized by Sigma (Table S3). The substrates were 5′-end labeled using T4 polynucleotide kinase and gamma-^32^P-ATP. For the label transfer reaction without competitors, recombinant SUMO-CFI_m_25 was first treated with protease Ulp1 (LifeSensors, Cat#SP4010) for one hour at 30°C (∼7 units of protease per 100 μg of SUMO-CFI_m_25). Cleaved or untreated SUMO-CFI_m_25 was then used in 10 μ L binding reactions at a final concentration of 3 μ M with 2 nM of the radiolabeled RNA substrates as well as 0.75% polyvinyl alcohol (PVA), 15 mM Tris-HCl pH 8, 15 mM NaCl, 25 mM KCl, 5 mM DTT, 11% glycerol, 10 mM HEPES, 0.1 mM EDTA, 0.25 μg *E. coli* tRNA, and 10 units RNasin. The reactions were performed with incubated at 30°C for 15 min. For the competition assays, 10 uL binding reactions were M SUMO-CFI_m_25 and 2 nM radiolabeled WT/UGUU RNA with the indicated concentrations of cold RNA competitor that had been treated at 70°C for 5 min then placed on ice. The reaction also included 0.75% PVA, 15 mM Tris-HCl pH 8, 40 mM KCl, 5 mM DTT, 7.5% glycerol, 1 mM HEPES, 0.01 mM EDTA, 0.25 μg *E. coli* tRNA, and 10 units RNasin. The reaction was incubated at 30°C for 30 min. All samples were crosslinked at 860 mJ/cm^2^ on ice ∼2 cm from a 254 nm UV light source (Spectroline XL-1500). The samples were resolved by SDS-page, the gel was dried and then bands analyzed by Phosphorimager.

#### Plasmids

Plasmids pX458-MAT2A-E9 and pX459-MAT2A-E8 are derived from pSpCas9(BB)-2A-GFP (pX458) and pSpCas9(BB)-2A-Puro (pX459) V2.0 which were gifts from Dr. Feng Zhang (Addgene plasmids #48138 and #62988). The targeting sequence was inserted using BbsI digestion and annealed 5′ phosphorylated oligonucleotides NC2980 and NC2981 (Table S3; pX458-MAT2A-E9) or NC2978 and NC2979 (Table S3; pX459-ΔMAT2A-E8) as previously described (Table S3)(Ran et al., 2013). To make pBS-ΔRI-Donor, we amplified the left homology arm with primers NC2982 and NC2983 (Table S3) and the right homology arm with primers NC2984 and NC2985 (Table S3) using cDNA from methionine-starved cells as a template to ensure PCR products had no intron 8. We used SOEing to join these products (Horton, 1995) and then inserted them into pBluescript II SK+ using NotI and BamHI restriction sites.

MAT2A β-globin reporter variants were made using β-MAT-WT from Pendleton et al., 2017. For the UGUU to UG**C**U mutation between the stop codon and hp1, β-MAT-WT was digested with XbaI before two PCR fragments amplified by NC3289 and NC3290 or NC3287 and NC3288 were inserted via Gibson Assembly Cloning kit (New England Biolabs)(Table S3). The UGUU to UGU**<u>A</u>** mutation was likewise made by Gibson assembly. First, NC2935 and NC1747 or NC3700 and NC3346 were used to amplify β-MAT-WT. Then the two PCR products were inserted via Gibson assembly into β-MAT digested with EcoRI and XhoI. The DI mutations (UG**C**A and **C**GUA) were made in a manner similarly to the UGUU to UGU**<u>A</u>** mutation. Two PCR products (see Table S3 for digested with EcoRI and XhoI. The DI mutations (UG**C**A and **C**GUA) were made in a manner similarly to the UGUU to UGU**A** mutation. Two PCR products (see Table S3 for primers) were amplified from β-MAT-WT then inserted into β-MAT-WT digested with EcoRI and XhoI via Gibson assembly. Mutations to the UGUA motifs found in the MAT2A 3′UTR were produced by amplifying a synthesized DNA fragment (GeneWiz) with NC3667 and NC3668 then insertion by Gibson assembly into β-MAT-WT digested with ApaI.

A plasmid containing the CFI_m_25 cDNA was obtained from the McDermott Sanger Sequencing Core. CFI_m_25 was amplified by NC3604 and NC3752 and then inserted into the pcDNA-flag and pcMS2-NLS-flag vectors using BamHI and XhoI sites. Similarly, CFI_m_25 derivatives were produced using SOEing using NC3604 and NC3752 as the exterior flanking primers and inserted into the same vectors using BamHI and XhoI sites. The internal primers for mutagenesis are listed in Table S3.

CFI_m_68 and CFI_m_59 were amplified as two fragments each from cDNA produced by oligo-dT priming, using primers listed in Table S3. The fragments were joined by SOEing using the forward primer from PCR#1 and reverse primer from PCR#2 then ligated into pcMS2-NLS-flag using BamHI and XhoI sites. Constructs with the RS domain deleted were produced by amplification from the pcMS2-NLS-fl-CFI_m_68 or - CFI_m_59 vectors using primers listed in Table S3 before insertion into pcMS2-NLS-flag using BamHI and XhoI sites.

pLentiV2-MAT2A was produced by annealing NC2533 and NC2534 before insertion into LentiCRISPRv2 (Addgene #52961; gift of Dr. Feng Zhang) following the Zhang lab protocol (Ran et al., 2013; Sanjana et al., 2014; Shalem et al., 2014). pLentiV2-NT was produced similarly, by annealing NC3198 and NC3199 before insertion into LentiCRISPRv2.

The pE-SUMO-CFI_m_25 expression plasmid was generated by amplification of CFI_m_25 cDNA with NC3852 and NC3853. The PCR product was digested with BbsI and XbaI and ligated into pE-SUMO cut with BsaI.

The construct pcbD1-MAT-E8-3′ HP1AG, (2xMS2) was generated by SOEing joining amplicons generated with primers NC1576 and NC2671 with those from SP6+ and NC2670. PCR reactions used pcβD1-MAT-E8-3′ HP1AG (No MS2) plasmid as a template. The products were inserted into β-MAT-WT cut with EcoRI and XhoI. AAVS1 1L TALEN and AAVS1 1R TALEN were gifts from Dr. Feng Zhang (Addgene plasmids #35431 and #35432)(Sajana et al., 2014). The plasmid hAAVS1-GFP-β2-MAT-E8-3′ was generated in two steps. First, we made pcGFP-β1-MAT-E8-3′ by Gibson assembly of three DNA fragments using the Gibson assembly. One insert was generated by amplifying EGFP using primer pair NC2229/NC2230 with pEGFP-N1 as a template and the second insert was made with primer pair NC2231/NC2232 using β-MAT-WT as a template. The vector fragment was generated by gel purification of β-MAT-WT cut with HindIII. The resulting plasmid was used as a template for PCR amplification with primers NC2264/NC2265 and the product was inserted into pAAVS-EGFP-DONOR digested with FseI and XbaI by Gibson assembly to generate hAAVS1-GFP-β2-MATE8-3′.

### Quantification and statistical analysis

Imagequant 5.2 was used to quantify northern blots. Bands were boxed at equal sizes in respective columns, with the rolling ball method used to subtract background. Image Studio Ver 3.1 was used to quantify western blots. Bands were selected using “Add Rectangle” feature with background automatically subtracted.

CRISPR screen data was analyzed by MAGeCK (see CRISPR screen methods)(Li et al., 2014). Poly(A)-ClickSeq was analyzed by ClickSeq Technologies using DPAC (Elrod et al., 2019; Routh, 2019; Routh et al., 2017). Venn diagrams were analyzed by SuperExactTest in RStudio (Wang et al., 2015). All other statistical analysis performed used two-tailed, unpaired student’s t-tests, with mean and standard deviation displayed. When p-value not listed or ns = not significant, *p ≤ 0.05, **p ≤ 0.01, ***p ≤ 0.001.

## Data and Code Availability

The Poly(A)-ClickSeq data has been deposited to the NCBI GEO database under the accession number GEO: GSE158591.

## Supplemental Figure Titles and Legends

**Figure S1.**
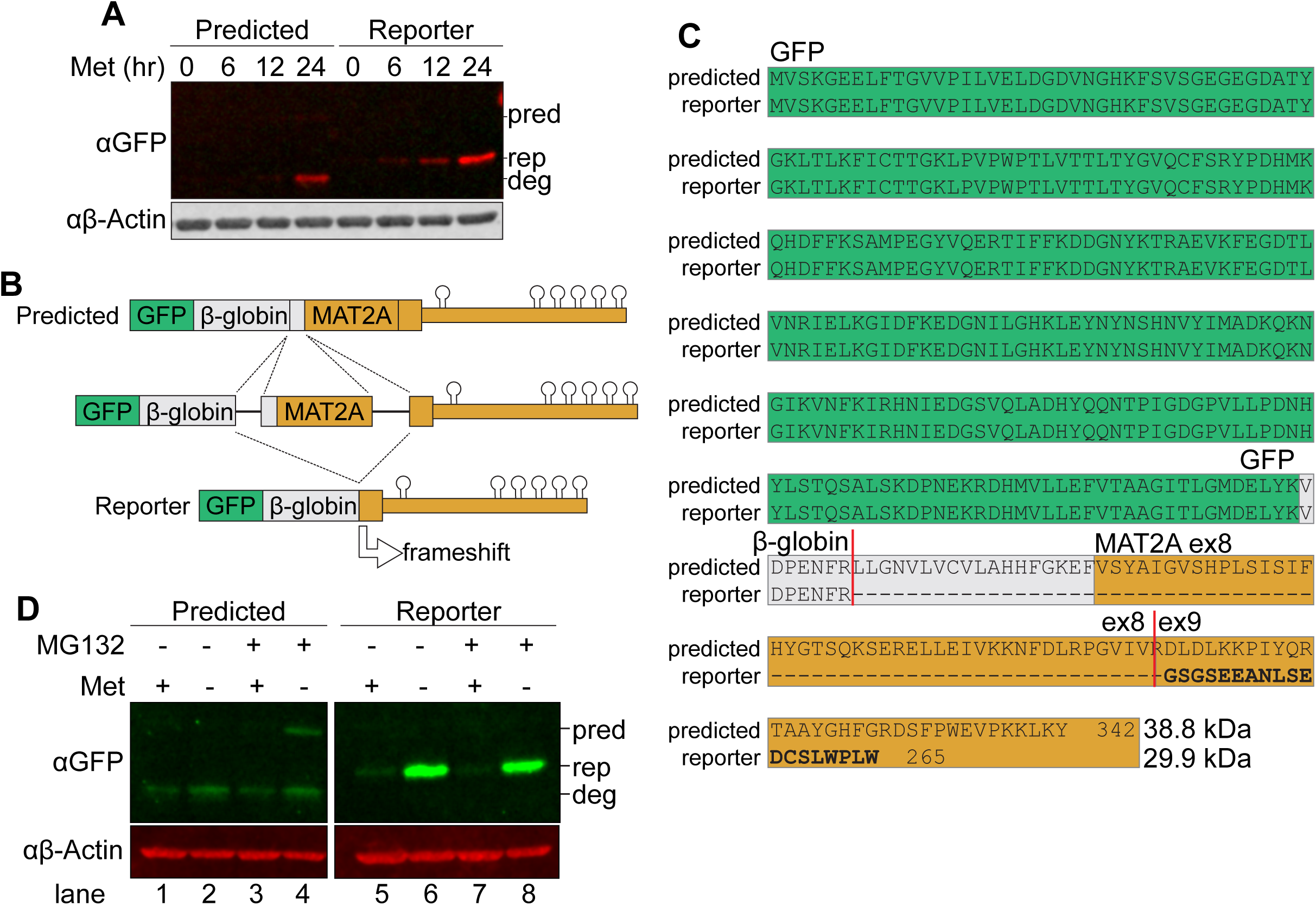
**An Alternatively Spliced Reporter Provides a more Robust GFP Signal, Related to Figure 1** A. Western analysis of GFP expression in two clonal reporter cell lines, the “Predicted” line splices in the expected pattern and the “Reporter” line is the one we used in our CRISPR screen. The “predicted” cell line appeared to have two bands, one faint band at the predicted MW (pred) and a second band of higher motility which is possibly a degradation product (deg). The reporter cell line had a single band at the size corresponding to the MW predicted from the alternative splicing pattern (rep). Actin serves as a loading control. B. Diagram of GFP reporter mRNA splicing. Splicing from the β-globin 5′ splice site to the MAT2A 3′splice site in our SAM-responsive clonal cell line results in a frameshift producing a smaller fusion protein. Diagram not to scale. C. Sequence alignment of proteins produced from the predicted and observed splicing of the reporter cell line. Green, GFP. Gray, β-globin. Orange, MAT2A exon 8-9. Predicted splice junctions shown by vertical red lines. D. Western analysis of GFP protein in a cell line with predicted splicing pattern or the reporter after 12 hr methionine depletion and 6 hr MG132 treatment. Images cropped from the same blot at same exposure. pred, predicted GFP-fusion protein size from predicted splicing pattern. rep, GFP product produced from reporter cell line. deg, putative degradation product from predicted splicing pattern product.

**Figure S2.**
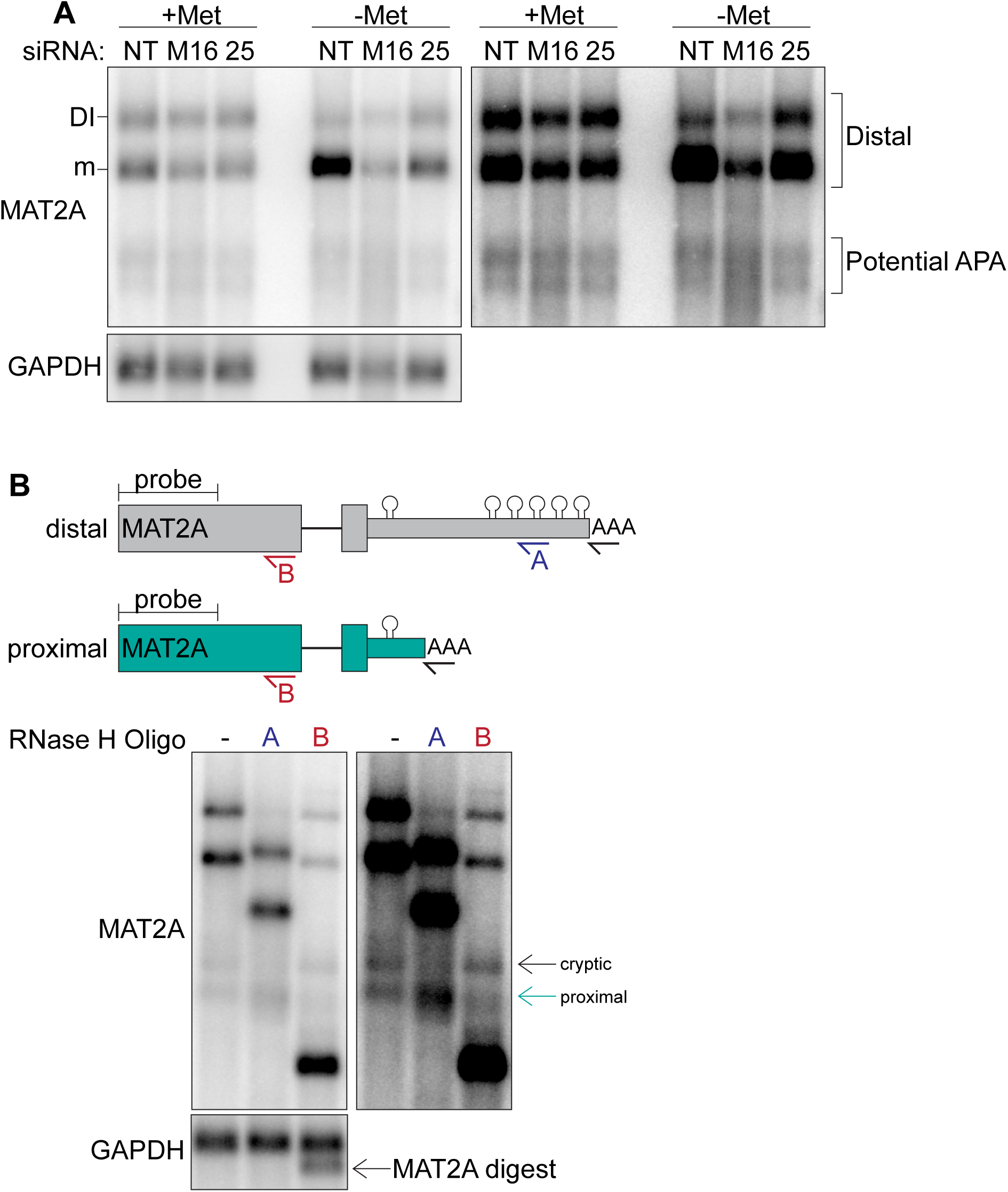
**MAT2A has undetectable APA regulation in response to methionine or CFI_m_25 in 293A-TOA cells, related to Figure 2.** A. Poly(A) selected RNAs were used for northern analysis of MAT2A isoforms after knockdown with non-targeting (NT), METTL16 (M16), or CFI_m_25 (25) siRNAs. Prior to harvesting, cells were conditioned in methionine-rich or -free media for four hrs. The blot on the right is overexposed image of the left. Distal refers to the distal poly(A) site used by the full-length MAT2A DI and mRNA isoforms. B. RNase H mapping of MAT2A isoforms using poly(A) selected RNAs and detected by northern analysis. *Top,* diagrams including DNA oligonucleotide positions used to target RNase H; both predicted distal and proximal isoforms are shown (not to scale). All samples were treated with dT_20_ (black) to create sharper bands by removing poly(A) heterogeneity. Oligonucleotide A (blue) is in MAT2A 3′UTR downstream of the predicted proximal poly(A) site, so it should not digest the proximally terminated RNA. In contrast, oligonucleotide B (red) hybridizes immediately upstream of the MAT2A exon 7-8 junction, so it should cleave both proximal and distal transcripts. Since B did not cleave the upper isoform, we labeled it “cryptic” due to its unknown origin. However, the pattern of the band marked “proximal” is consistent with usage of the proximal poly(A) site.

**Figure S3.**
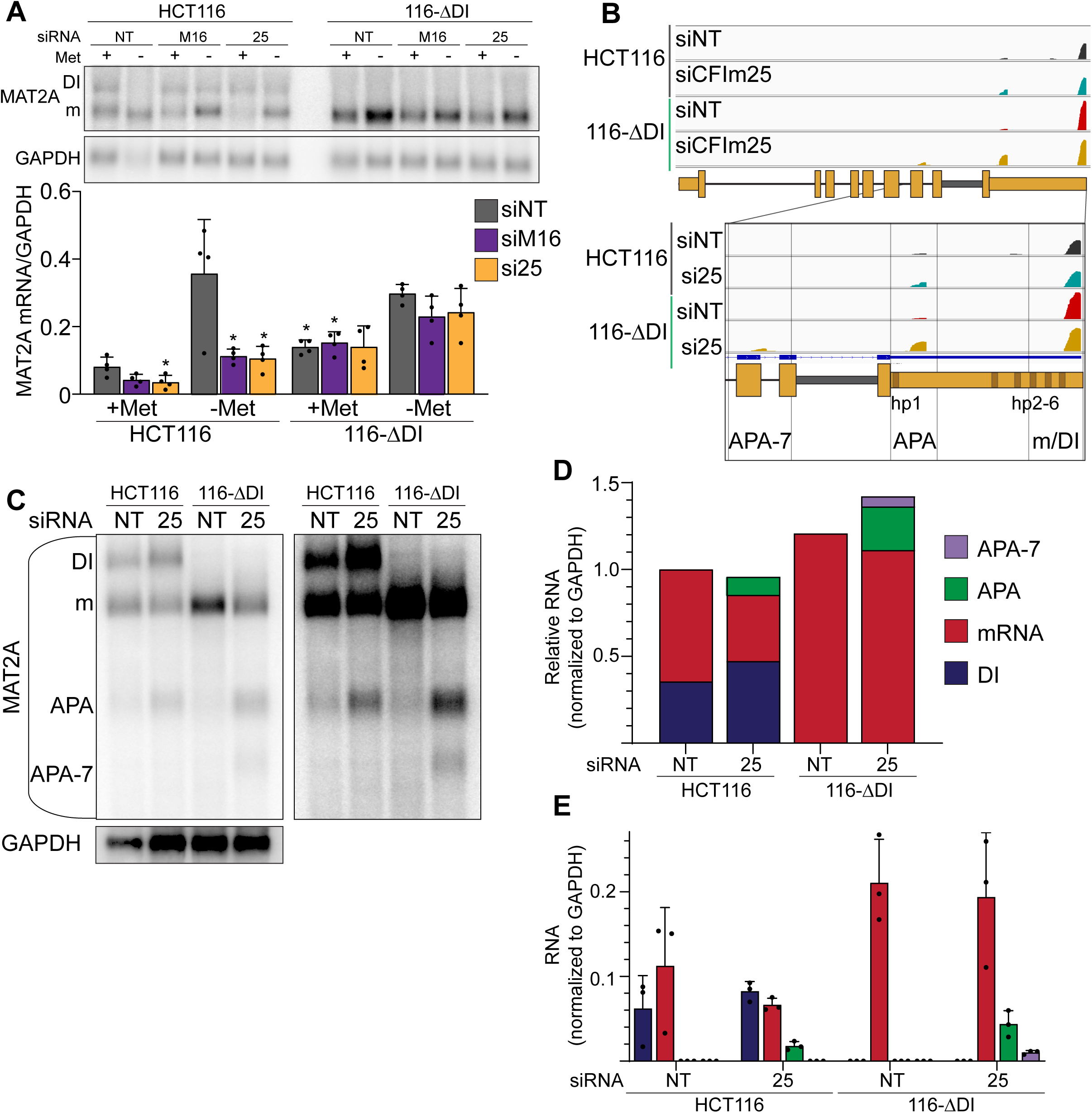
**MAT2A isoforms in HCT116 and 116-**Δ**DI, Related to Figure 3** A. Northern blot analyses of MAT2A expression in HCT116 and 116-ΔDI cells upon knockdown of the indicated factor after four hours methionine depletion where indicated. Asterisks denote significance relative to HCT116 siNT sample under the same methionine conditioning. n = 4. B. PAC-seq trace for MAT2A. The magnified region is from exon 7 to the end of the 3′UTR. Black and blue, HCT116 siNT and siCFI_m_25, respectively. Red and yellow, 116-ΔDI siNT and siCFI_m_25, respectively. APA-7, alternative poly(A) site found near the beginning of intron 8. APA, proximal poly(A) site found between hp1 and 2. m/DI, distal poly(A) site used in prevalent mRNA and DI isoforms. One representative sequencing trace shown of three biological replicates. C-E. Representative northern blot (C) and quantification (D, E) of MAT2A isoforms after CFI_m_25 knockdown and poly(A) selection in HCT116 and 116-Δ DI cell lines. Quantification shown as stacked (D) or individually with error bars (E). Blue, DI. Red, mRNA. Green, APA site found in the 3′UTR. Purple, APA-7, the APA site found within intron 7. n = 3.

**Figure S4.**
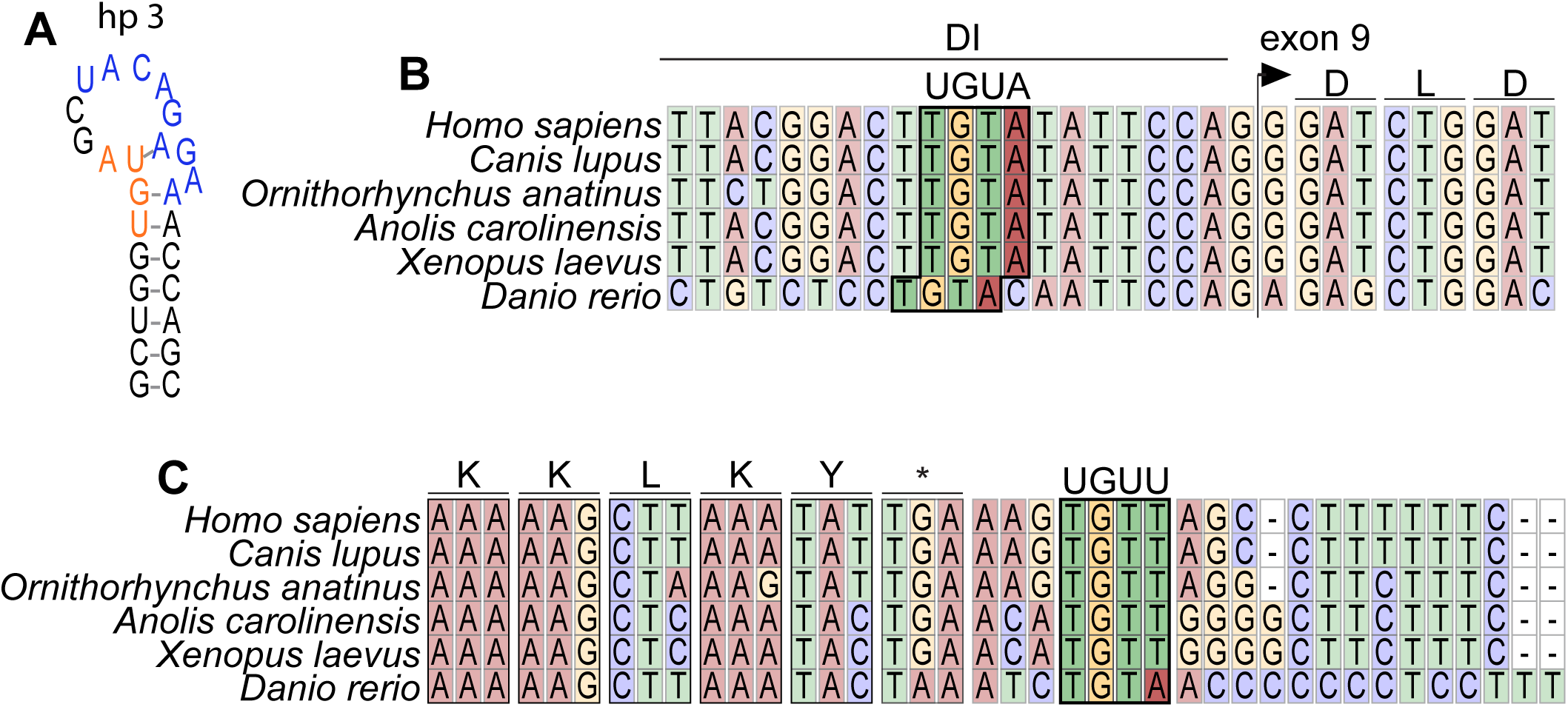
**MAT2A contains conserved CFI_m_25 binding sites in the detained intron and 3′UTR, Related to Figure 4** A. MAT2A hp3 diagram modeled on crystallographic data of hairpin 1 (Doxtader et al., 2018). A UGUA site (orange) is embedded in critical structural nucleotides for hairpin folding and recognition by METTL16. Blue, METTL16 binding motif. The upstream GU is also conserved in METTL16 binding sites to generate the appropriate RNA structure, but only hp3 has the GU within a UGUA motif. B. Vertebrate sequence alignment of MAT2A surrounding the predicted UGUA binding motif in the detained intron (boxed). C. Vertebrate sequence alignment of MAT2A surrounding the predicted UGUU binding motif (boxed).

**Figure S5.**
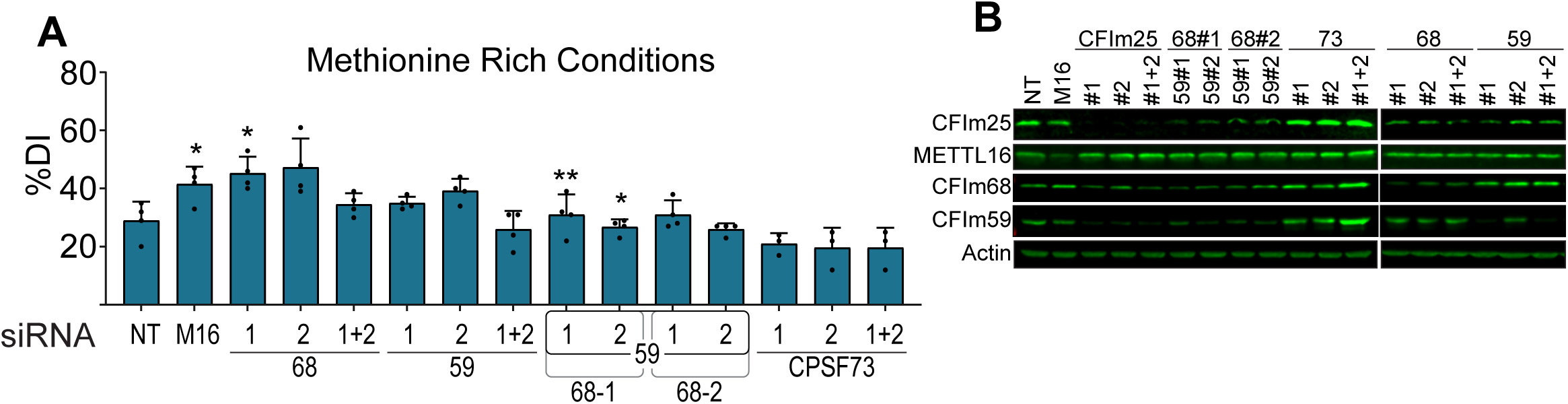
**Additional controls for the CFI_m_ complex knockdown experiments, Related to Figure 5** A. Quantification of MAT2A intron detention after CFI_m_68 and CFI_m_59 depletion in methionine-rich conditions. Two independent siRNAs for each factor were used (labeled #1 and #2). CPSF73 knockdown serves as a negative control. Matched representative northern blot and methionine-depleted quantification in 5A and B. n ≥ 3. B. Western analysis of protein expression after knocking down the indicated factors individually or in combination. The first five lanes are the same blot as Figure 2C, but here we include additional knock-down conditions.

**Figure S6.**
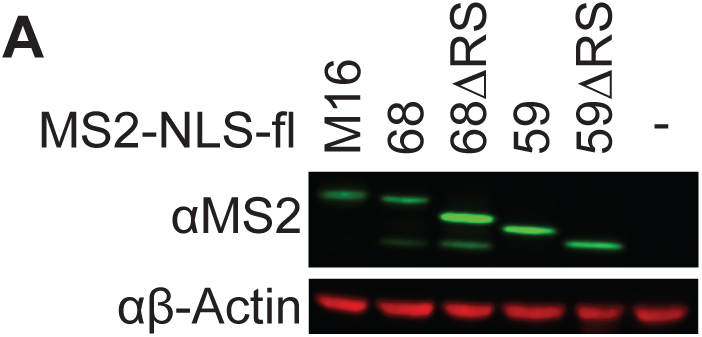
**Expression of full-length and RS-deletion MS2-NLS-fl CFI_m_68 and CFI_m_59 proteins. Related to Figure 6.** A. Western analysis of protein expression for MS2-NLS-fl tagged METTL16, CFI_m_68 and CFI_m_59 using anti-MS2 antibodies. Expression of the 17 kDa MS2-NLS-fl has been confirmed, but it is too small to be detected under these conditions.

**Figure S7.**
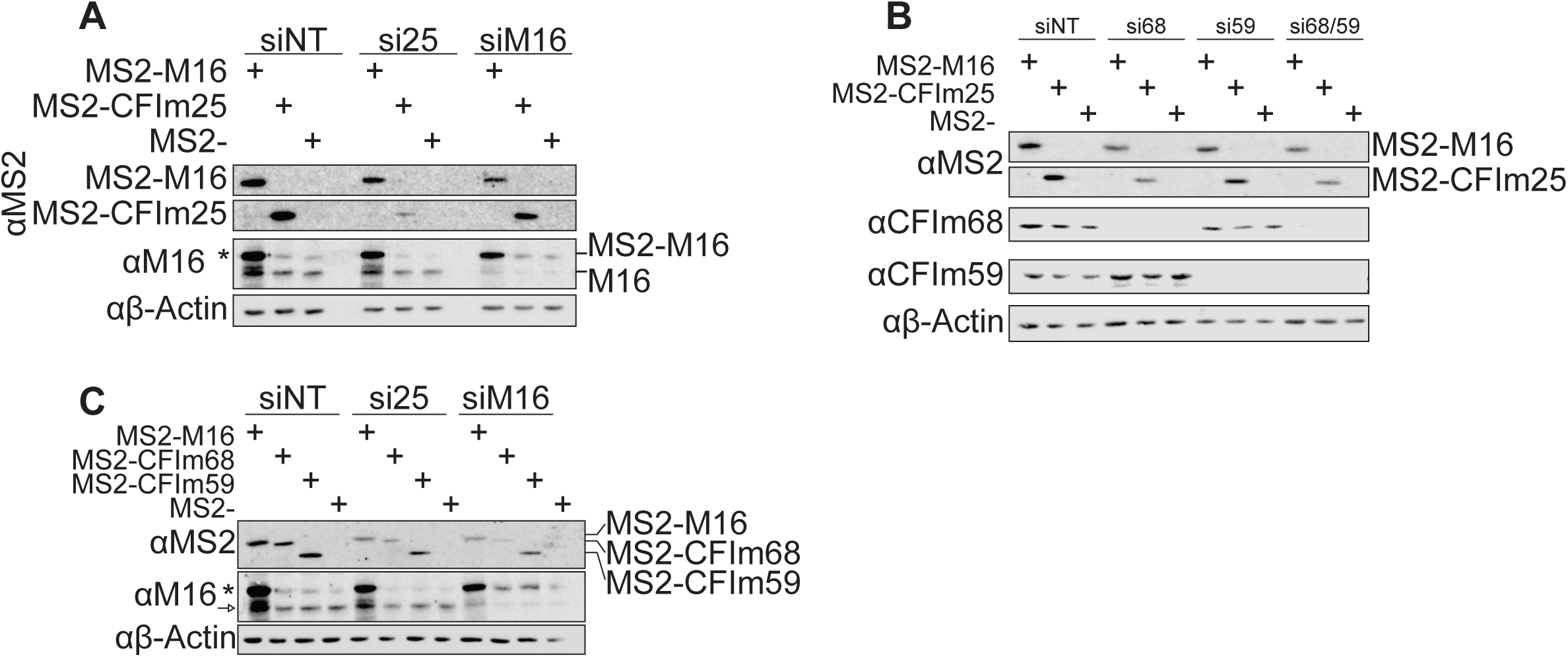
**Expression of MS2-NLS-fl proteins after depletion of METTL16 or CFI_m_ components. Related to Figure 7** A. Western analysis of MS2-NLS-fl tagged METTL16 and CFI_m_25 RNA binding mutant after knocking down the indicated protein. The METTL16 construct is siRNA-resistant to siM16, while the CFI_m_25 construct is not resistant to siCFI_m_25. Asterisk denotes MS2-NLS-fl tagged METTL16 detected by METTL16 antibody. Related to Figure 7B. B. Western analysis of MS2-NLS-fl tagged METTL16 and CFI_m_25 RNA binding mutant after CFI_m_68/59 depletion. Related to Figure 7C. C. Western analysis of MS2-NLS-fl tagged METTL16, CFI_m_68, and CFI_m_59 after METTL16 or CFI_m_25 depletion. Asterisk denotes MS2-NLS-fl tagged METTL16 detected by METTL16 antibody. Arrow denotes endogenous METTL16. Related to Figure 7D.

In some cases, the knockdown of these essential factors lowers the expression of the MS2 transgenes (Fig S7). However, it is important to note that the reduced levels of MS2-CFI_m_25 observed upon METTL16 knockdown potently drive splicing (Fig 7B, purple). Conversely, the levels of MS2-NLS-fl-METTL16 observed upon CFI_m_25 knockdown is reduced, but these levels are the same to those observed upon METTL16 knockdown and they remain well above the endogenous levels (Fig S7C). In this case, these levels of expression also are sufficient to drive splicing (Fig 7D, purple). Therefore, we conclude that the changes in transgene expression are not responsible for the observed changes in β-globin-MAT2A reporter splicing.

## Excel Table Titles and Legends

**Table S1. CRISPR screen results**

MAGeCK analysis of biological replicates of CRISPR screen. Table shows MAGeCK analysis for each individual replicate (rep1-3) and analysis of the replicates together (triplicate). The triplicate analysis is referenced in text and Figure 1.

**Table S2. Poly(A)-ClickSeq results**

DI cell lines with either siNon-targeting Poly(A)-ClickSeq analysis of HCT116 and 116-Δ or siCFI_m_25 treatment.

**Table S3. Nucleic acid reagents**

DNA oligonucleotide sequences for cloning primers, northern blotting probe primers, RNase H oligonucleotides, CRISPR screen primers, and synthesized DNA for cloning. RNA oligonucleotides sequences for label transfer assay substrates.

## KEY RESOURCES TABLE

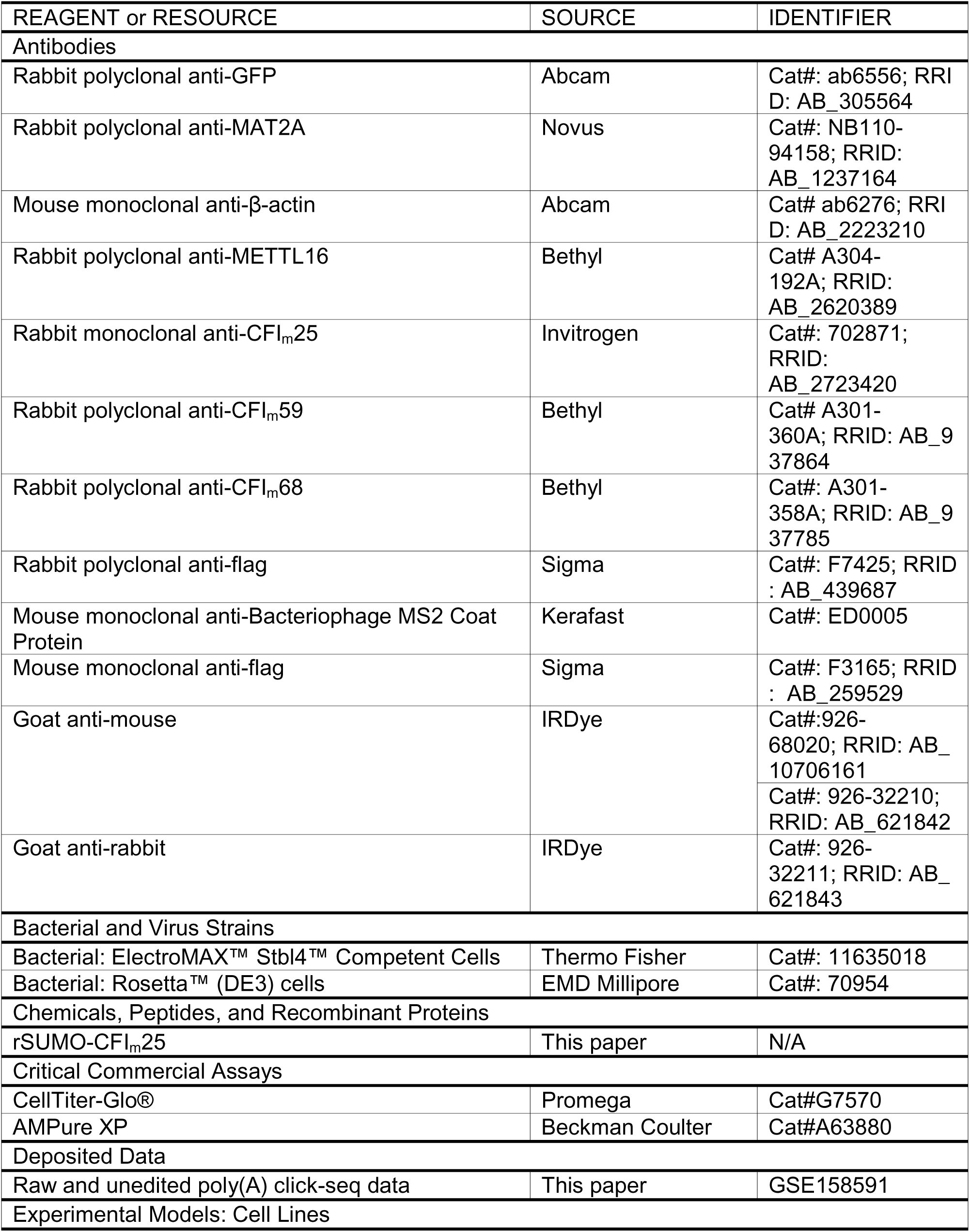
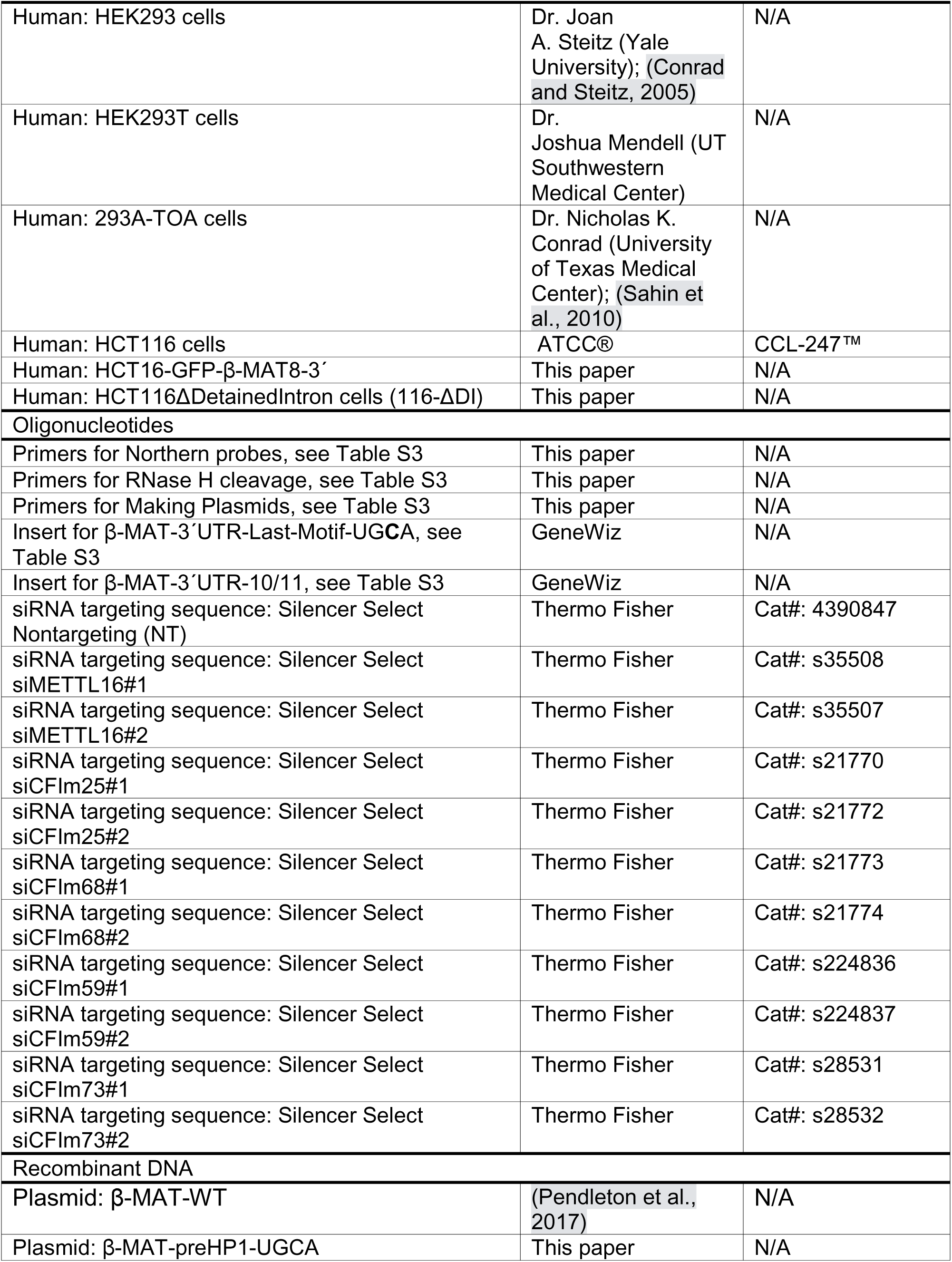
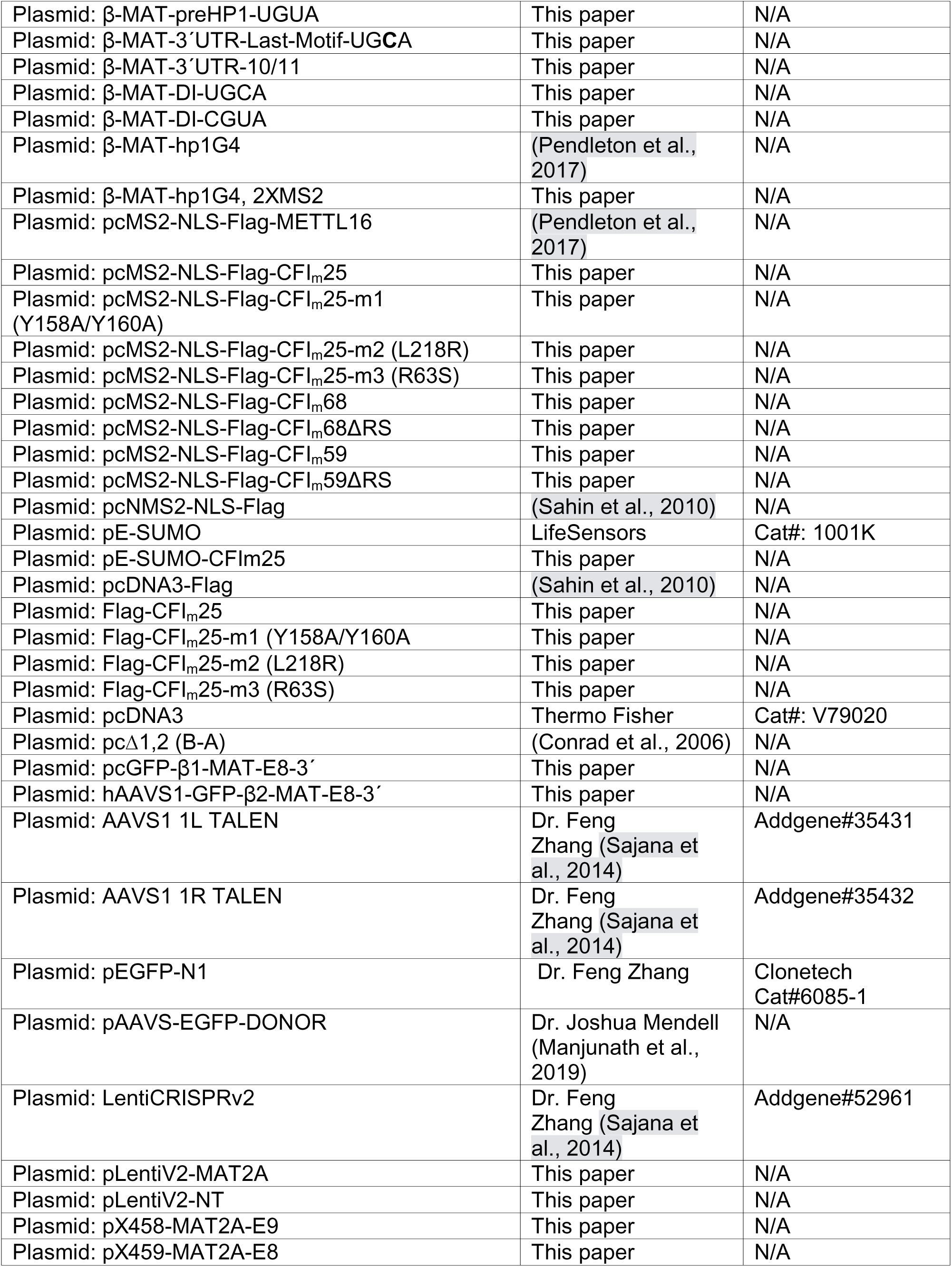
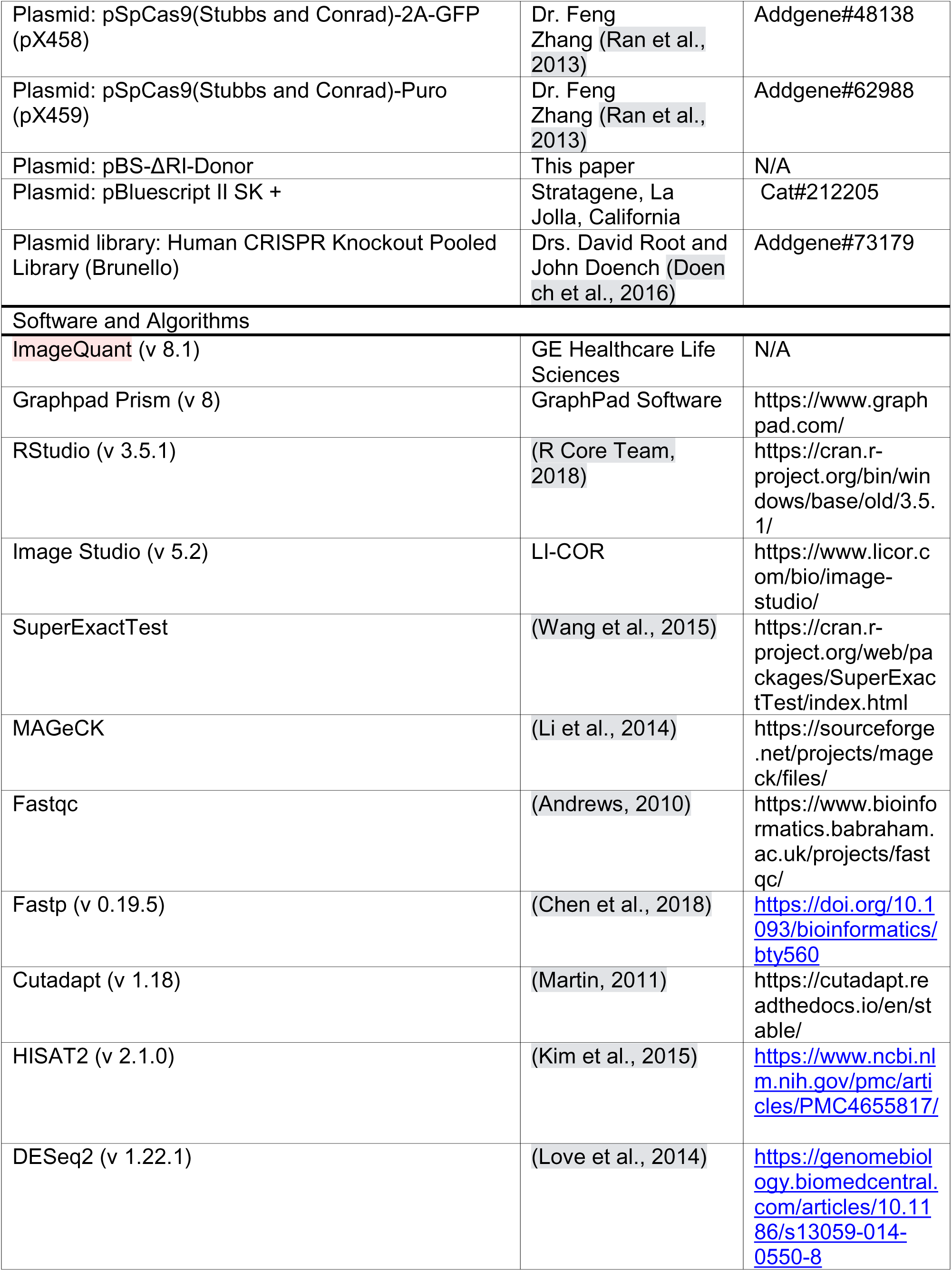
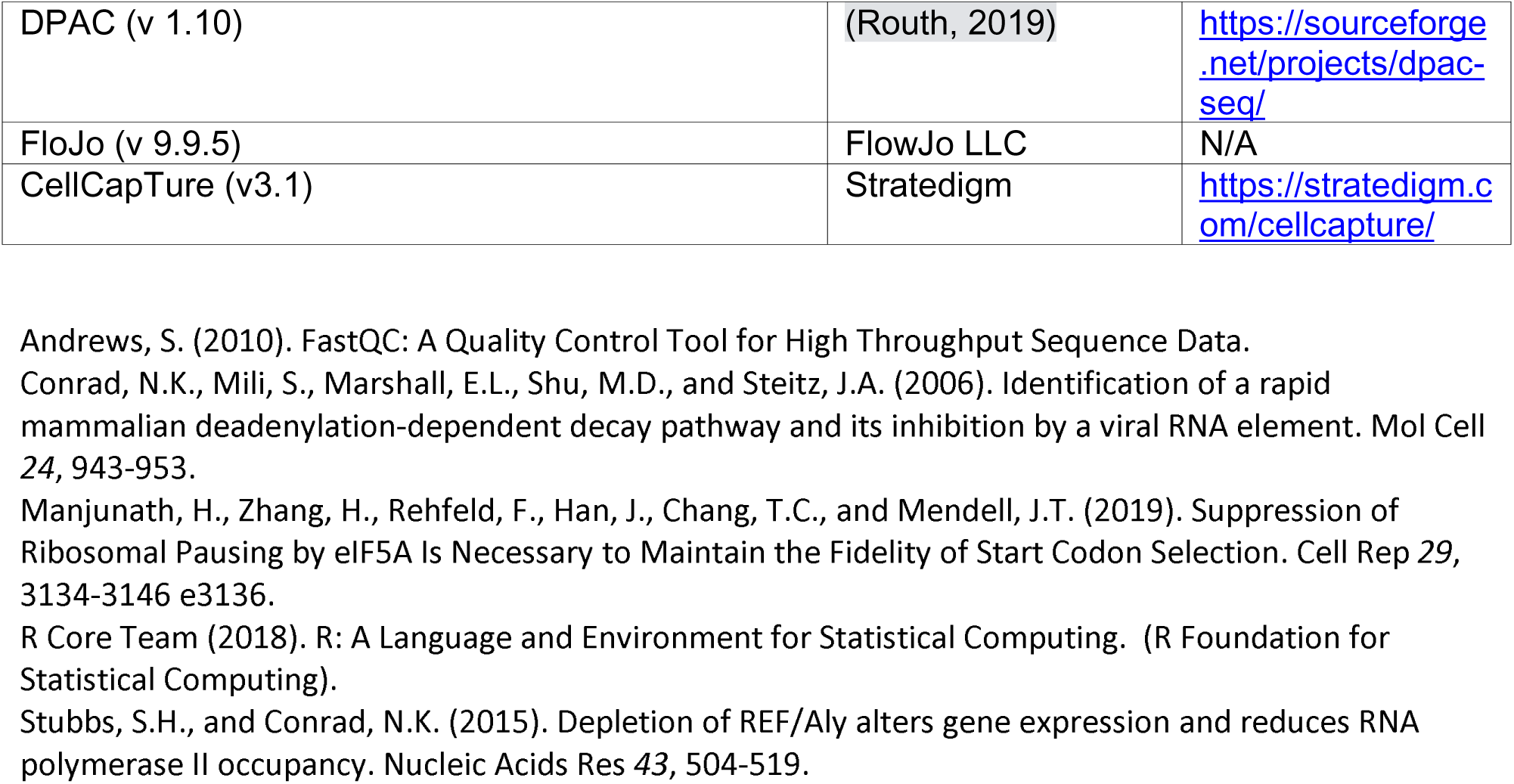

